# An emergent temporal basis set robustly supports cerebellar time-series learning

**DOI:** 10.1101/2022.01.06.475265

**Authors:** Jesse I. Gilmer, Michael A. Farries, Zachary Kilpatrick, Ioannis Delis, Abigail L. Person

## Abstract

Learning plays a key role in the function of many neural circuits. The cerebellum is considered a ‘learning machine’ essential for time interval estimation underlying motor coordination and other behaviors. Theoretical work has proposed that the cerebellum’s input recipient structure, the granule cell layer (GCL), performs pattern separation of inputs that facilitates learning in Purkinje cells (P-cells). However, the relationship between input reformatting and learning outcomes has remained debated, with roles emphasized for pattern separation features from sparsification to decorrelation. We took a novel approach by training a minimalist model of the cerebellar cortex to learn complex time-series data from naturalistic inputs, in contrast to traditional classification tasks. The model robustly produced temporal basis sets from naturalistic inputs, and the resultant GCL output supported learning of temporally complex target functions. Learning favored surprisingly dense granule cell activity, yet the key statistical features in GCL population activity that drove learning differed from those seen previously for classification tasks. Moreover, different cerebellar tasks were supported by diverse pattern separation features that matched the demands of the tasks. These findings advance testable hypotheses for mechanisms of temporal basis set formation and predict that population statistics of granule cell activity may differ across cerebellar regions to support distinct behaviors.

## Introduction

The cerebellum refines movement and maintains calibrated sensorimotor transformations by learning to predict outcomes of behaviors through error-based feedback (Ito, 1972; Herzfeld et al., 2015; Medina 2000; Mauk and Buonomano, 2004; Raymond et al., 1996). A major site of cerebellar learning is in the cerebellar cortex, where Purkinje cells (P-cells) receive sensorimotor information from parallel fibers (Huang et al. 2013) whose synaptic strengths are modified by the conjunction of presynaptic (parallel fiber) activity and climbing fiber inputs to P-cells thought to convey instructive feedback (McCormick et al., 1982; Yang and Lisberger, 2014; Mauk et al., 1986; De Zeeuw et al., 1998). P-cell activity is characterized by rich temporal dynamics during movements, representing putative computations of internal models of the body and the physics of the environment (Wolpert et al., 1998; Shadmehr and Mussa-Ivaldi 1994). Parallel fibers are the axons of cerebellar granule cells (GCs), a huge neuronal population (comprising roughly half of the neurons in the entire brain; Herculano-Houzel 2010), which are the major recipient of extrinsic inputs to the cerebellum. Thus, understanding the output of the GCL is key in determining the encoding capacity and information load of incoming activity projected to the cerebellum. Inputs to GCs arise from mossy fibers (MFs), which convey sensorimotor information used by the cerebellum to predict the consequences of motor commands (Rancz et al., 2007; Ishikawa et al., 2015). There are massively more GCs than MFs and each GC typically receives input from just 4 MFs (Palkovits et al., 1971), such that the information carried by each MF is spread among many GCs but each GC samples from only a tiny fraction of total MFs (Jakab and Hamori 1988; Eccles et al., 1967).

The GCL has been the focus of theoretical work spanning decades that has explored the computational advantages of the unique architecture of the structure. Notably, early studies of the cerebellar circuit by Marr (1969) and Albus (1971) proposed that a key component of the cerebellar algorithm is the sparse representation of MF inputs by GCs. In this view, the cerebellum often must discriminate between overlapping, highly correlated patterns of MF activity with only subtle differences distinguishing them (Bengsston and Jorntell 2009). Sparse recoding of MF activity in a much larger population of GCs (“expansion recoding”) increases the dimensionality of population representation and transforms correlated MF activity into independent activity patterns among a subset of GCs (Litwin-Kumar et al., 2017; Cayco-Gajic et al., 2017; Gilmer and Person 2018). These decorrelated activity patterns are easier to distinguish by learning algorithms operating in P-cells, leading to better associative learning and credit assignment (Cayco-Gajic et al., 2017; Sanger et al., 2020).

The machine learning perspective of Marr-Albus theory tends to assume that the cerebellum is presented with a series of static input patterns that must be distinguished and categorized. However, neuronal population dynamics are hardly ever static and precise timing of circuit inputs to the cerebellum remains an essential part of cerebellar function. Mauk and Buonomano (2004) revisited cerebellar expansion recoding in the context of delayed eyeblink conditioning, a cerebellum-dependent learning task where the subject hears a tone followed by an aversive air puff to the eye at a fixed delay from tone onset and must learn to initiate an eyeblink at the correct delay to protect the eye. They proposed that a static activity pattern in MFs (representing the tone) could be recoded in the GC layer as a temporally evolving set of distinct activity patterns. P-cells could learn to recognize the GC activity pattern present at the correct delay and initiate an eyeblink to avert the “error” signal representing the air puff to the eye. In other words, P-cells would select from a “temporal basis set” for correct error prediction and learning adaptive behavior.

Expansion recoding creates the possibility of representing a single MF pattern as a series of distinct GC patterns (a “temporal basis set”; Albus 1975; Zhou et al., 2020; Tyrrell and Willshaw 1992; Liu et al., 2019; Kalmbach et al., 2011). The existence of this predicted temporal basis set within the cerebellum has been supported experimentally in electric fish, where GCs represent the duration of mimicked electric organ discharge through a range of onsets (Kennedy et al., 2014). Although these studies have been highly influential, little is known about how the GCL would produce a temporally diverse basis set from static input data. Local inhibition, short-term synaptic plasticity, and varying GC excitability all may work together to diversify time-invariant input (Chabrol et al., 2015; Duguid et al, 2012; Crowley et al., 2009; Rudolph et al., 2015; Buonomano and Mauk 1994; Kanichay and Silver 2008; Simat et al., 2007; Mapelli et al., 2009; Rossi et al., 1996; Gall et al., 2005; Armano et al., 2000; Rizwan et al. 2016; Tabuchi et al., 2019; D’Angelo and De Zeeuw 2009). However, the assumption that MFs ever provide truly static input to the cerebellum is probably unrealistic; even a static stimulus like a tone will generate time-varying activity patterns in the auditory brainstem as units undergo adaptation (Eriksson and Robert 1999). Moreover, most of the input signals that the cerebellum must process are intrinsically dynamic (Bengsston and Jorntell 2009; Chabrol et al., 2015). We seek to explore how expansion recoding of dynamic, naturalistic input activity assists cerebellar function.

To test how expansion recoding of naturalistic input contributes to learning, we developed a simple model of the GCL and a time-series prediction task to explore the effect of putative GCL filtering mechanisms on expansion recoding and learning (Fig. 1A). Similar to previous models, this simplified model made GC activity sparser relative to MF inputs (Marr 1969; Albus 1971) and increased the dimensionality of the input activity (Litwin-Kumar et al., 2017) while preserving information (Billings et al., 2014). That these features of GCL function were achieved using only basic approximations of GC physiology suggests that the crystalline connectivity and feedforward inhibition of the cerebellum incorporated in our model are sufficient to produce pattern separation of naturalistic time-varying inputs. This model demonstrates greatly enhanced learning accuracy and speed by P-cells on a difficult time series prediction task when compared to MF inputs alone. Although we observed robust sparsening of input activity by GCL output, the relationship between pattern separation metrics and the observed learning was dependent upon the task being performed, suggesting that GCL output covers a span of modalities supporting flexible feature selection by P-cells to meet the needs of particular learning targets. These findings reinforce the ideas explored previously that the GCL balances input sparsening against information loss to optimize learning (Cayco-Gajic et al., 2017; Cayco-Gajic and Silver 2019), and that the balance between these features of GCL output can be functionally controlled through adjustments in the strength of local inhibition. We conclude by showing that muscle activity during reaching movements (Delis, et al. 2018), a proxy for time-varying efference copy signals received by the cerebellum, gives rise to information-preserving sparseness that supports time-series predictions, suggesting that physiological input sources to the GCL, like the spinocerebellar pathways, are sufficient to drive learning. Together, these results suggest that the cerebellar GCL provides a rich basis for learning in downstream Purkinje cells, providing a mixture of lossless representation (Billings et al., 2014) and enhanced spatiotemporal representation (Litwin-Kumar et al. 2017) that are selected for by associative learning to support the learning of diverse outputs that support adaptive outputs in a variety of tasks (Fujita 1982; Dean and Porrill 2008).

**Figure 1:**
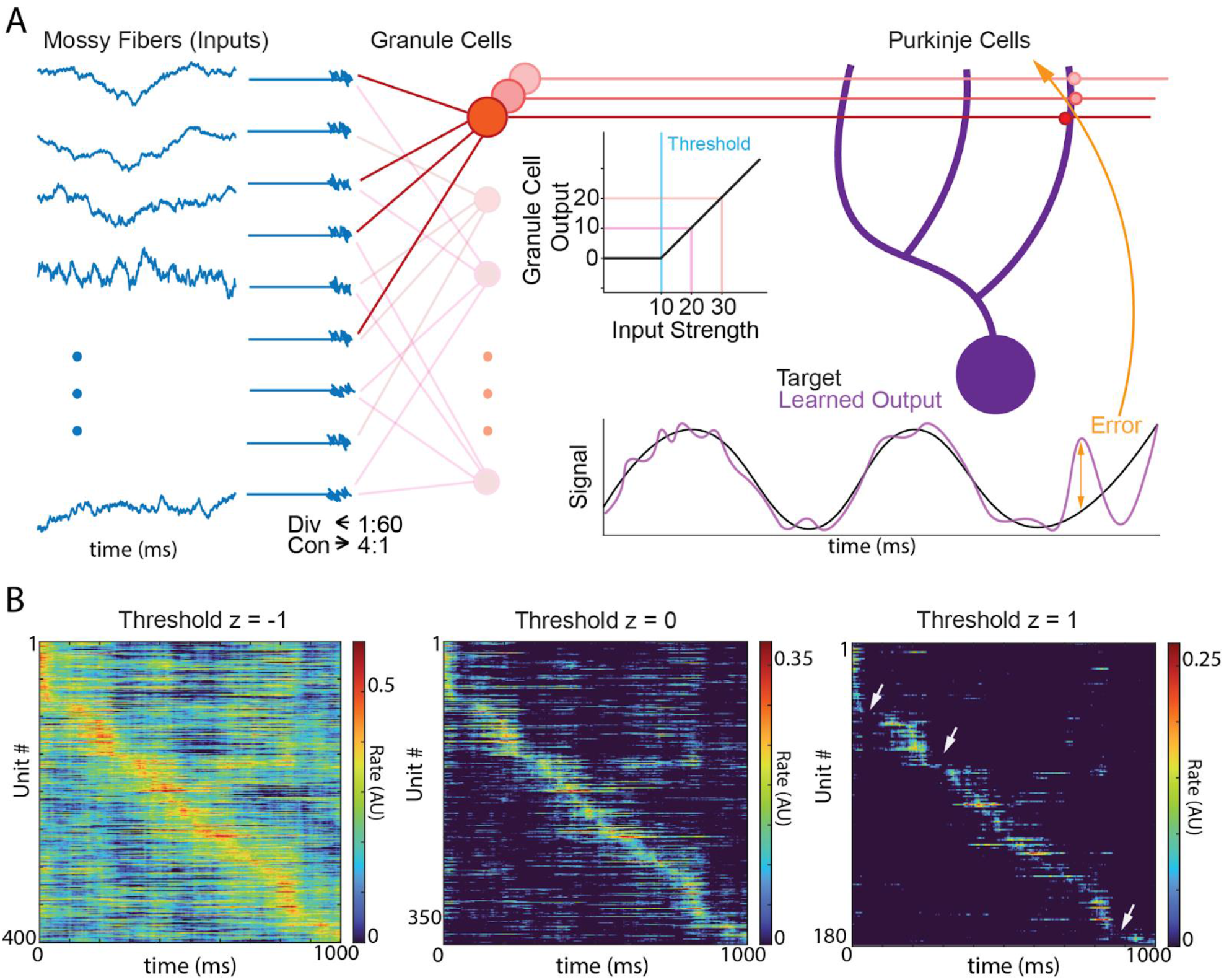
Model architecture and effects of thresholding on GCL population activity. **A**. Diagram of algorithm implmentation. Left shows Ornstein-Uhlenbeck processes (see Methods) as proxies for mossy fiber (MFs, blue) inputs to granule cell (GCs, red) units, with convergence and divergence of MFs to GCs noted beneath MFs. GCs employ threshold-linear filtering shown beneath the red parallel fibers. GC outputs are then transmitted to downstream Purkinje cells (P-cells). P-cells learn to predict target functions based on summation of weighted GC inputs and differences between the prediction and true target are transmitted as an ‘error’, which determines the updates to the weights between GCs and P-cells. **B**. Example unit GC population rates when threshold is -1.0, 0 and 1.0 showing the gradual sparsening of GCL output. Arrows on 1.0 plot indicate regions of gaps in representation (lossiness) by the GCL population due to over-sparsening.

## Results

### Temporal basis set formation as emergent property of GCL filtering of physiological-like inputs

The cerebellar granule cell layer (GCL) is theorized to convert spatiotemporally dense inputs into discrete representations through coincidence detection and feedforward and feedback inhibition-mediated thresholding (Marr 1969; Solinas et al., 2010). How the GCL expands spatiotemporal representation has been the subject of debate and scientific inquiry for decades. While cellular and circuit mechanisms have been proposed to expand time invariant signals such as tones (Mauk and Buonomano 2004; Medina 2000), naturalistic cerebellar inputs are typically time varying by virtue of dynamic sensorimotor interactions with the environment (Rancz et al., 2007; Eriksson and Robert 1999). Moreover, cerebellar learning is thought to sculpt complex time-varying outputs in Purkinje cells (P-cells) that reflect behavioral adaptations. This observation raises the question of how GCL output supports time series learning, a divergence from traditional classification tasks used in cerebellar models. To address this, we investigated how such naturalistic input patterns were transformed by the GCL to support learning time-varying output patterns, such as those required for generating and correcting movements, or for producing predictions of sensory events (Fig. 1; Izawa et al. 2012).

We created a simple model capturing the dominant circuit features of the GCL: sparse sampling of mossy fiber (MFs) inputs by postsynaptic granule cells (GCs) and coincidence detection regulated by cellular excitability and local feedforward inhibition (Figure 1A; Eq.1,2; Marr 1969; Albus 1971; Palkovits et al., 1971; Chabrol et al., 2015). MF inputs are represented as smooth time-varying functions, i.e., as variable firing rates rather than spike trains. GC output is generated by summing MF inputs and thresholding the resultant sum; anything below threshold is set to zero while suprathreshold summed activity is passed on (minus the threshold) as GC output (Fig. 1A, center). The GC threshold level represents both intrinsic excitability and the effect of local feedforward inhibition on regulating GC activity. To model MF activity patterns, we sought a statistical ensemble that was rich enough to capture the dynamic nature of naturalistic inputs while remaining analytically tractable and easily parameterized. We chose to utilize the Ornstein-Uhlenbeck (OU) stochastic process, whose output is Gaussian and varies over an adjustable timescale. The statistics of an OU process can be fully characterized by just three parameters: mean, standard deviation, and correlation time; samples drawn from an OU process are shown in Fig. 1A (left, blue). Since the input to GCs is Gaussian in our model, the summed activity that is thresholded is Gaussian as well. For that reason, we found it convenient to define the GC threshold in terms of z-scores. Thus a GC with a threshold of “zero” would have its threshold set at the mean value of its MF inputs; such a GC would be silent 50% of the time on average because the Gaussian presynaptic input would be below the mean value half the time. This makes it possible to discuss functionally similar thresholds across varying network architectures (e.g., a GC with a threshold of zero would discard half of its input on average regardless of whether it received 2 or 8 MF inputs). Via this simple mechanism, our model GCL generates temporally sparse activity that could support learning by downstream P-cells (Fig. 1A, right). Indeed, when subjected to this form of filtering, the resultant representation in the GCL population became spatiotemporally distinct at moderate thresholding levels (near 0, Fig. 1B, center). However, too little thresholding resulted in dense representation (Fig. 1B, left) while too much thresholding resulted in over-sparsening, leading to loss of representation in the temporal domain (Fig. 1B, right, arrows indicate loss of representation). The emergence of sparse spatiotemporal representation under the simplistic constraints of the model suggests that the cerebellum’s intrinsic circuitry is sufficient to produce spatiotemporal separation when given sufficiently time-varying inputs.

**Figure 1, figure supplement 1:**
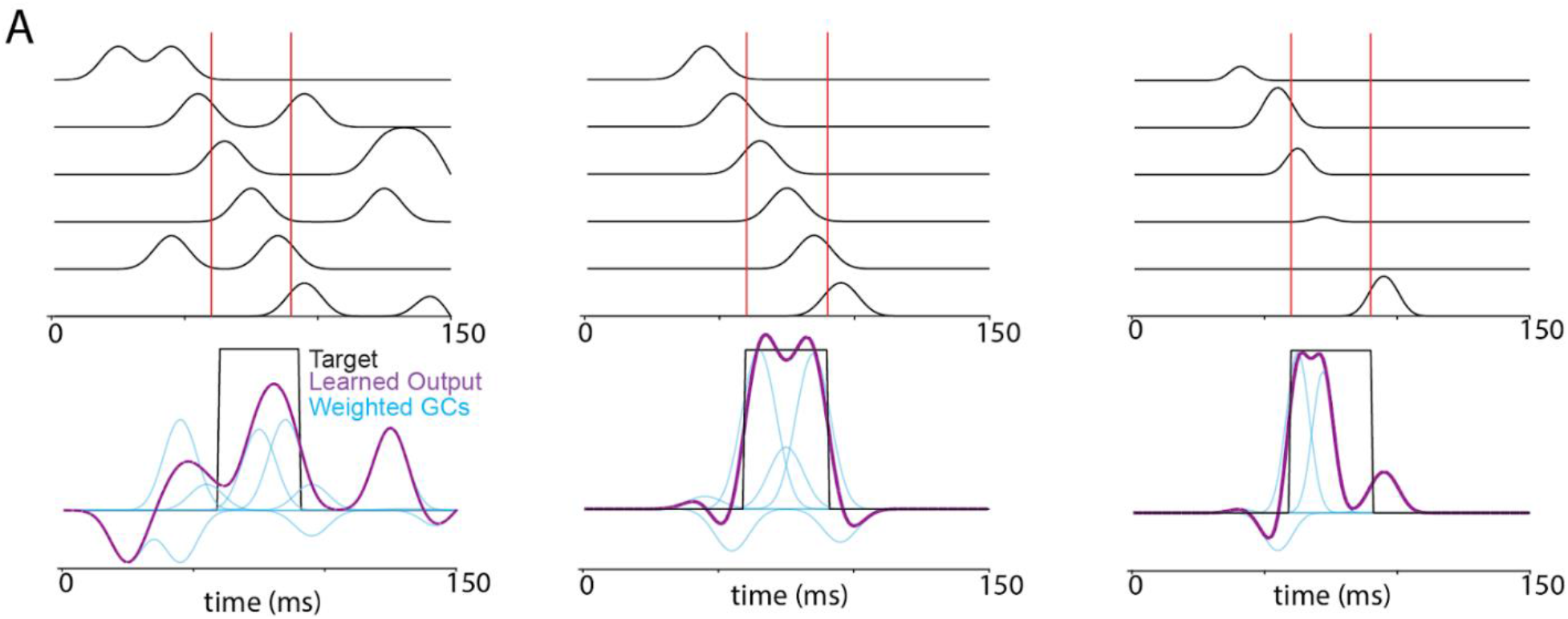
Example of basis set utility in learning. **A**. Diagram relating fictive GC activity (top) with resultant learning (bottom) using those fictive signals as the basis for learning. Target functions are shown in black and learned outputs with minimized error are shown in purple. Note that the best learning occurs with uniform, minimally overlapping GCs, tiling the epoch when the target signal is active (red lines; middle panel).

### GCL improves time series learning accuracy

Next, we tested whether GCL population activity seen above assisted learning. We devised a learning task where P-cells learned to generate a specific time-varying activity pattern in response to the dynamic activity patterns generated by MFs, which better represents the tasks performed by the cerebellum than pattern classification. The target patterns that P-cells were tasked with generating were drawn from an OU process with an autocorrelation time of 10 ms (see Methods). P-cells initially produced output very unlike the target, but over repeated trials their output converged towards the target function (see Fig. 3A for example progression of learning). We compared this convergence of P-cell output to target when input activity was filtered through the GCL to performance the case when MF activity is sent directly to P-cells (“MFs alone”). The GCL enhanced convergence to target at thresholds between –1 and 1 (Fig. 2A), achieving a minimized mean squared error (MSE) of roughly 0.005 compared to 0.02 when using MFs alone. It may seem that the performance with MFs alone was still quite good, if slightly quantitatively inferior, when compared to the range of the target function (normalized to a range of [0,1]). Thus, to establish intuition into the practical difference of this range of MSEs, we tasked the model with recapitulating a complex image with an identical range of target function values (with identical range of [0,1], Fig. 2B). Importantly, the model GCL generated a recognizable image, with an MSE of 0.002 while experiments using MF alone generated an unrecognizable image with an MSE of 0.02. (The relative MSE, i.e. the ratio of GCL MSE to MFs alone MSE, was 0.08). Thus, this MSE range represented the difference between noise and easily recognizable images and text (Fig. 2B top right vs three thresholds, bottom). This principle was qualitatively true of abstract target functions used in OU input experiments as well (Fig. 3A for example target functions and estimations). Thus, the inclusion of the GCL in the filtering process greatly improves learning of complex functions by P-cells in this task, supporting an order of magnitude improvement in MSE of learned target functions compared to MFs alone. Importantly, this was not a consequence of the large population expansion between MFs and GCs, as increasing the number of MFs alone did not improve performance to the levels observed in the model GCL (Supp. Fig. 2A), but a sufficiently large GCL population is required to improve learning (Supp. Fig. 2B).

**Figure 2:**
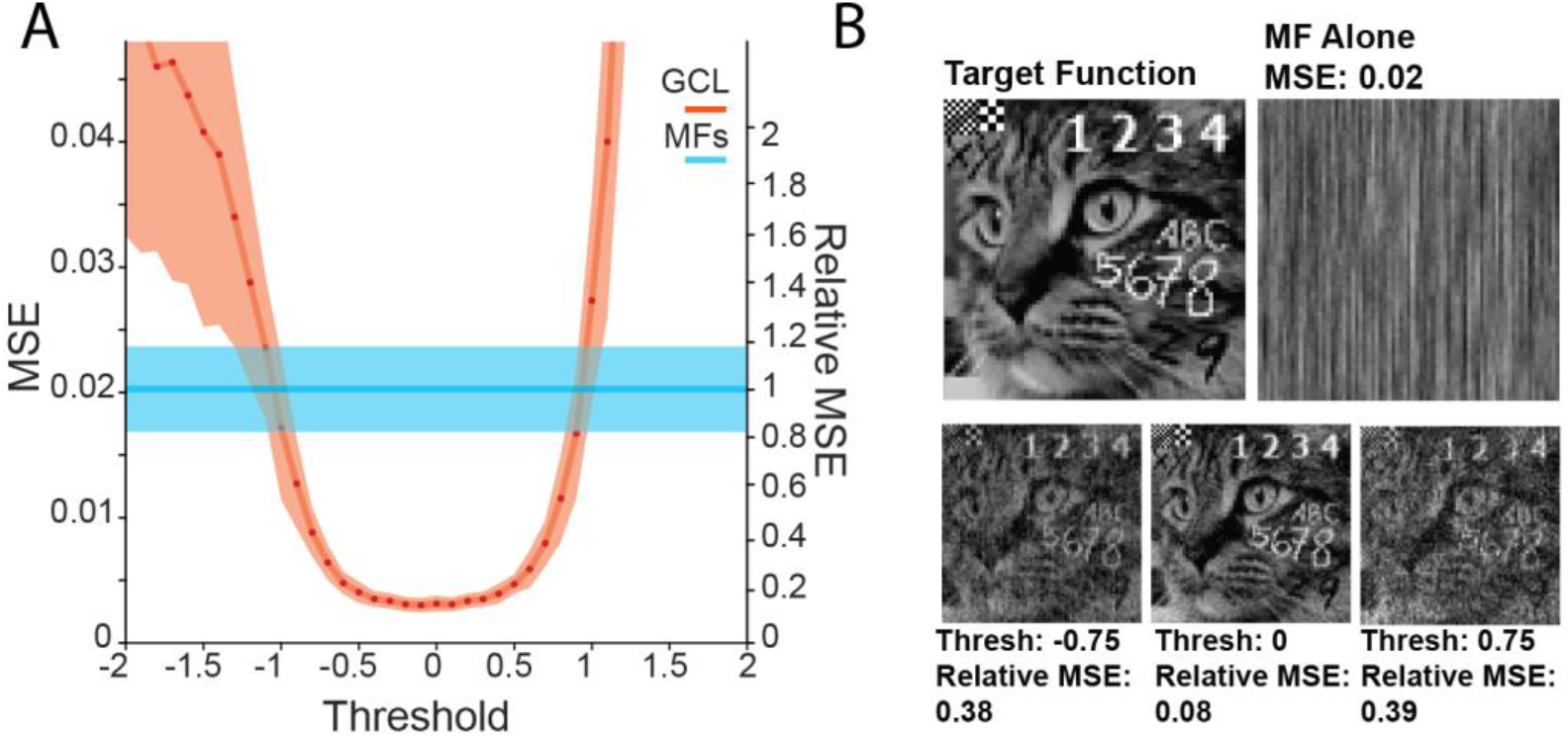
Enhanced time series learning using GCL model. **A**: Relation between mean squared error (MSE) and threshold in a 50 MF, 3000 GC system, showing a significant reduction in error between a threshold of –1 and 1 for the learning model using GCL output (orange) compared to mossy fibers alone (blue). Transparent bounds represent standard deviation of learning outcome. Relative MSE of the GCL is shown on the right margin and represents the ratio of MSE for the GCL compared to MF alone. Values less than 1 indicate GCL outperforming MFs alone. **B**. An intuitive demonstration of the difference in the small MSE change produced by the MF-direct task, and the much clearer MSE produced by the GCL model used as input to P-cells. Panels show the outcomes of the same task with the target function being an image of a cat, with both handwritten and typeface text, and a 1- and 2-pixel width checkerboard (upper left corner).

**Figure 3.**
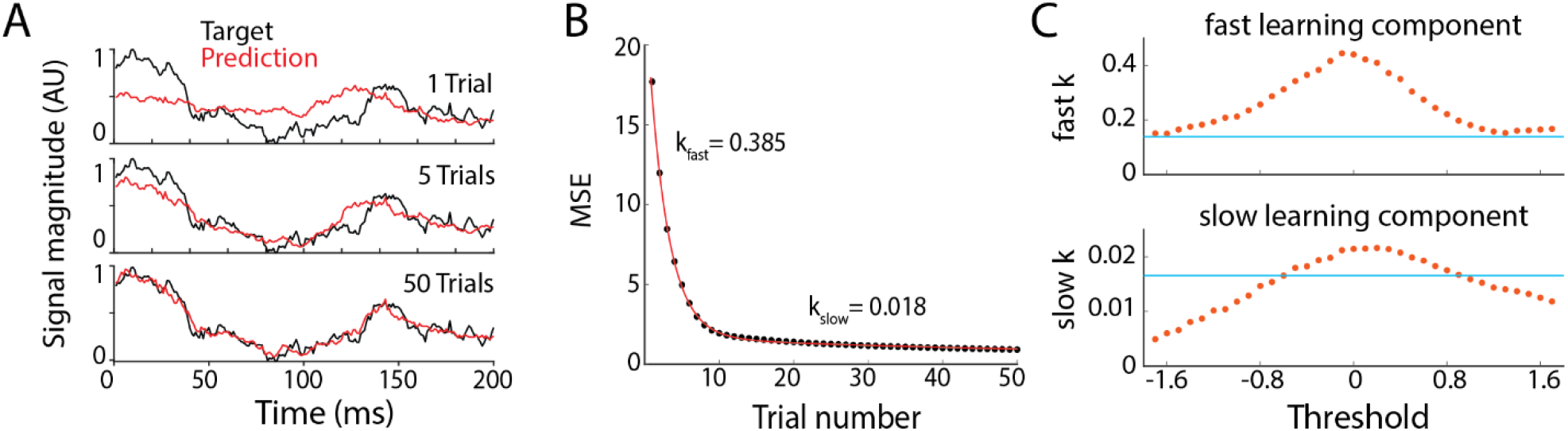
Learning speed increases with GCL. **A**. Example of learned predictions after 1,5, and 50 trials of learning, with predictions in red and target function in black. **B**. Example learning trajectory of MSE fit with a double exponential. Black circles: MSE of network output on each trial. Red line: double exponential fit MSE during learning. Here, step size was 10^−6^ and z-scored GCL threshold was 0. We use the exponents k from the exponential fit to measure learning speed. **C**. Learning speed as a function of GCL threshold (red dots). Blue line: learning speed in networks lacking GCL, i.e. mossy fibers directly innervate output Purkinje unit, gradient descent step size was 10^−6^.

**Figure 2, Figure Supplement 1:**
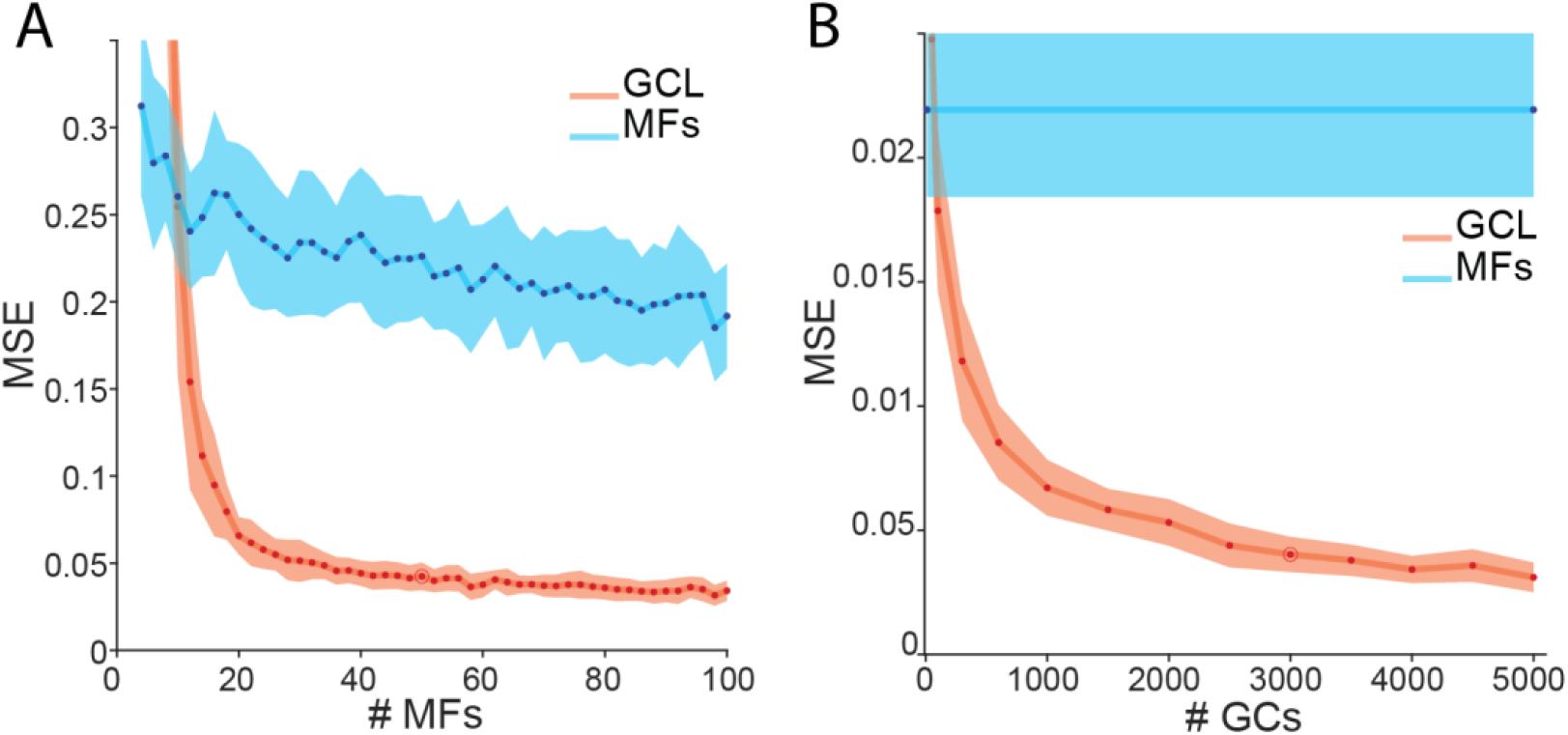
Effects of input and output population sizes on learning. **A**. Relation between the number of mossy fiber inputs and the resultant MSE, with MFs either inputted directly to P-cells (blue) or fed through 3000 GC unit model (red). **B**. MSE as a function of GC number compared to 50 MFs alone (blue). GC threshold fixed at 0 for these simulations.

### GCL model speeds time series learning

Having found that the GCL improves the match between predicted output and target output over a range of thresholds, we next examined whether the structure also increased the speed of convergence. Examining the MSE between output and target on each trial as training progresses (Fig. 3C, *red circles*), we found that output usually converged rapidly at first then more slowly in later stages of training (Fig. 3A). The reduction in MSE over training in our model was reasonably well fit by a double exponential (Fig. 3B, *red curve*), of the form

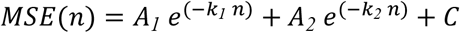

where *n* is the trial number. We measured the convergence speed of a simulation by the rate constants *k*_1_ and *k*_2_. In the vast majority cases, one of these rate constants was 5-50 times larger than the other; we denote the larger constant *k*_*fast*_ and the other *k*_*slow*_. For most parameter values, *k*_*fast*_ accounts for more than 80% of learning.

We next examined the influence of key parameters on convergence speed. First, we looked at the effect of the GC threshold. Learning was fastest for GCL thresholds near zero (Fig. 3C, *red circles*), the level that filters out half of the input received by a GC. Convergence in networks that lack a GCL (MFs directly innervating P-cells) was consistently slower (Fig. 3C, *blue line*) than networks with a GCL. Convergence can also be sped up by increasing the size of the parameter jumps in synaptic weight space during gradient descent (the “step size”), but only to a limited degree (Supp. Fig. 3A). Indeed, at a GCL threshold of 0, convergence speed *decreased* as the step size size was increased beyond ∼10^−6^ (au). We speculated that this trade-off was a consequence of a failure to converge in a subset of simulations. To test this, we looked at the fraction of simulations that converged towards a low MSE as a function of the update magnitude. We found that the fraction of simulations that converged (“fraction successful”) decreased with increasing step size (Supp. Fig. 3B); in simulations that did not converge, the MSE increased explosively and synaptic weights diverged. In such cases, we assume the large weight updates made it impossible to descend the MSE gradient; each network weight update drastically changed the cost function such that local MSE minima were overshot. When larger step sizes did permit convergence, progress was nevertheless slowed, likely because the relatively large learning rates led to inefficient progress towards the MSE minimum.

Although larger step sizes eventually cause learning to slow and then fail entirely at a given GCL threshold, higher thresholds permitted larger step sizes before failures predominated (Supp. Fig. 3B). Since higher thresholds permit larger step sizes before convergence failure sets in, convergence speed might be maximized by jointly optimizing step size and GCL threshold. We tested this by systematically raising step sizes at each threshold until convergence success fell to 50%. We defined the “maximum convergence rate” for a given threshold as the maximum convergence rate (derived from fitting the MSE trajectory with a double exponential) yielding successful convergence at least 50% of the time. We found that the threshold giving the fastest convergence was indeed higher when step size was also optimized (Supp. Fig. 3B) than when step size was fixed (Fig. 3C). Thus, increased GCL thresholding can allow the network to trade learning accuracy for increased speed of learning.

**Figure 3, figure supplement 1:**
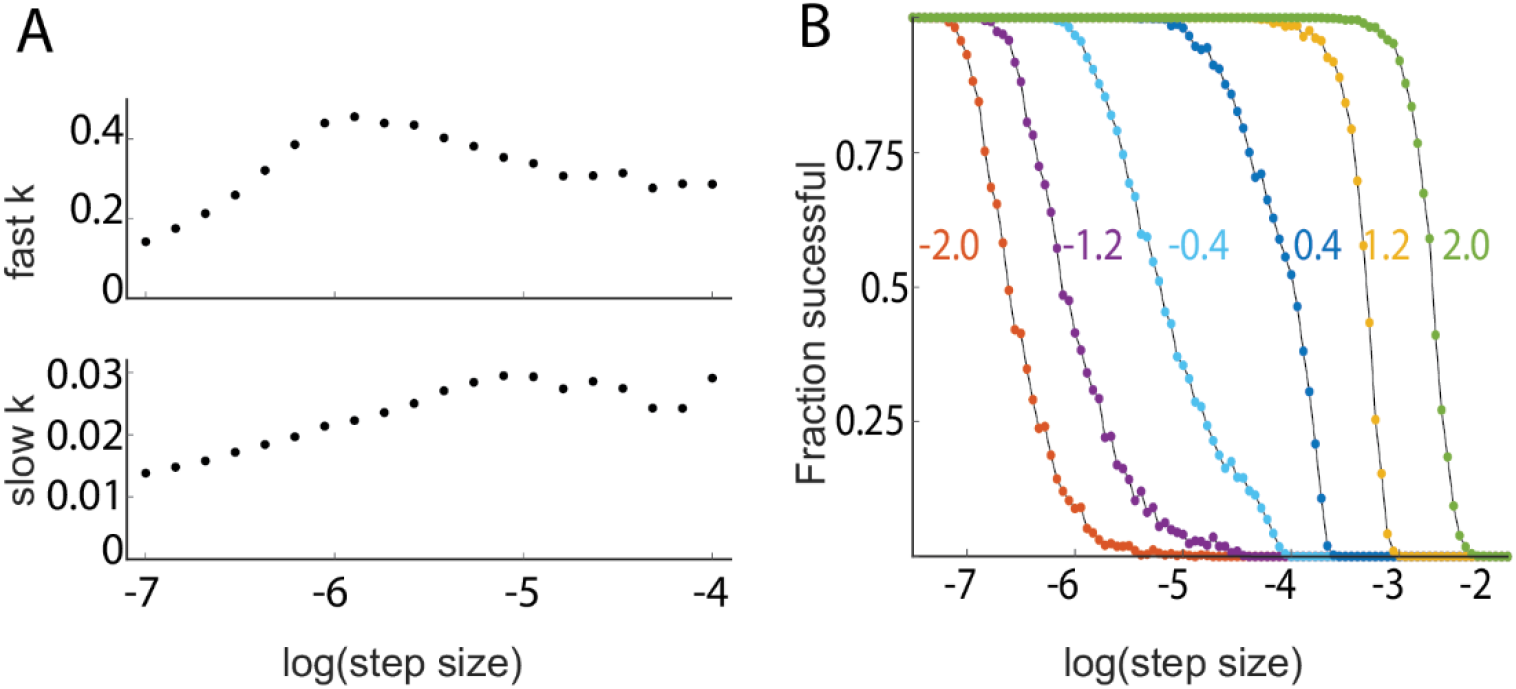
Effects of gradient descent step size on learning speed. **A**. Learning speeds (exponential time constant) for different simulations using varying gradient descent step sizes, showing differentially maximized learning speeds occurring at different step sizes. **B**. Fraction of simulations that converge to asymptotic MSE values as a function of gradient descent step sizes for different values of GCL threshold (colors denote threshold values). Note that larger step sizes and faster learning are supported in models with higher thresholds.

### Recovering GCL input from GCL output

Having established a framework for studying GCL processing of naturalistic inputs, we wanted to understand to what extent thresholding GCL activity led to the loss of information supplied by MF inputs, which potentially contains useful features for learning. In other words, would Purkinje neurons be deprived of behaviorally relevant mossy fiber information if these inputs are severely filtered by the GCL? To assess this issue, we used a metric of information preservation called *explained variance*, (Achen 1982); however, in this special case, we use the term ‘*variance retained*’, because this metric represents the preservation of information about the input after being subjected to filtering in the GCL layer. Let *x*_*t*_ denote the MF input at time *t*. If the GCL activity preserves the information present in *x*_*t*_, then it should be possible to reconstruct the activity of MFs from GCL activity (see Methods for details on how this reconstruction was performed). The variance retained is then the mean squared error between the actual MF input *x*_*t*_ and the reconstructed input, normalized by the MF input variance:

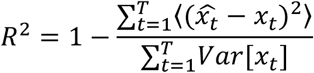

Our primary finding is that the GCL transmits nearly all of the information present in the MF inputs even at fairly high thresholds, but only if the GCL is sufficiently large relative to the MF population. The threshold, feedforward architecture, and relative balance of MF inputs and GC outputs all affect the quality of the reconstruction. Variance retained by the reconstruction layer decreased with the GC layer threshold, since it masked some subthreshold input values (Fig. 4B). Allowing more MF inputs per GC recovered some of this masked information, since some subthreshold values are revealed through summing with sufficiently suprathreshold values. However, these gains cease beyond a few MF inputs per GC, since the exponential growth of MF combinations rapidly exceeds the number that the GCs can represent (Marr 1969; Gilmer and Person 2017).

**Figure 4:**
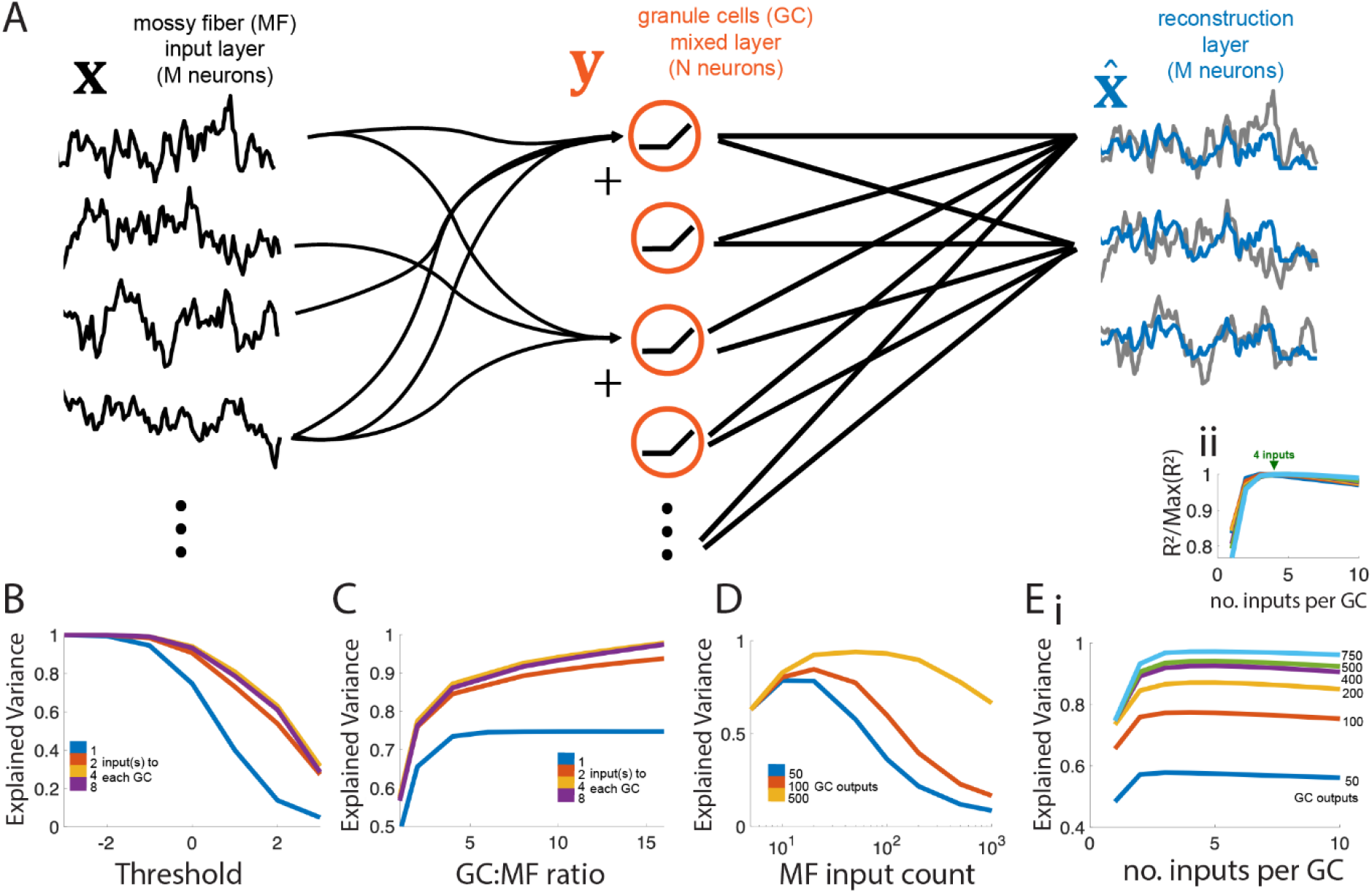
Recovering inputs with an optimal linear readout. **A**. Network model schematic. Granule cell (GC, red, center) layer thresholds the sum of (4 here) randomly chosen mossy fiber (MF, black, left) inputs, which are then fed into a reconstruction layer which uses the optimal weighting from all N GCs to approximate each of the M inputs (compare blue readouts to grey inputs). **B**. Increasing the threshold of the GC layer (N=500 outputs) decreases the explained variance (i.e. variance retained) of the best reconstruction layer (M=50), but the effect is reduced with an intermediate number of MF inputs per GC. **C**. Variance retained increases with the ratio of GCs per MF but gains from increasing the number of inputs to each GC are limited (max at 4 inputs). Here there are M=50 MF inputs at the threshold = 0. **D**. For a fixed number of GC outputs N, there is an optimal number of MF inputs (M) for which the variance retained of the reconstruction layer is maximized. **E. i**. For a fixed number of GC outputs N and MF inputs M=50, there is an optimal number of inputs per G (around 4) for maximizing variance retained. **ii**. Same as i, but with each value normalized to its maximum to show maximized values at inputs = 4.

To disentangle the information contained in the summed inputs, many different combinations of inputs must be represented to disambiguate the contributions of each MF input. Increasing the number of GCs generally increases the variance retained, since more combinations of MF inputs are represented, and reveal subthreshold input values (Fig. 4C). Interestingly, variance retained by the network varied non-monotonically with the number of MF inputs (M) when the number of GCs (N) was fixed. This is because having too few MF inputs means there may not be a sufficient number of combinations so that subthreshold values can be revealed (by summing them with suprathreshold inputs) but having too many saturates the information load of the GC layer (Fig. 4D). Lastly, when fixing the number of MF inputs and GCs, there is an optimal number of MF inputs to each GC, which aligns with the anatomical convergence factor of 4 MF/GC (Fig. 4E), related to previous findings that suggest the best way to maximize dimensionality in the GC output layer is to provide sparse input from the mossy fibers (Litwin-Kumar et al., 2017; Cayco-Gajic et al., 2017). Thus, there are two key features that shape the information transferred to the GCL from the MF inputs. First, the way in which MF inputs are combined to form the total input to each GC determines how much information about subthreshold inputs can be transferred through the nonlinearity. Second, the total number of GC outputs determines how many MF input combinations can be represented, so that, ultimately, the random sums of MFs can be disentangled by the downstream reconstruction layer. Together, information transfer requires a combined summation and downstream decorrelation process accomplished by the three layer network.

### General statistical features of GCL population activity

To better relate the present model to previous theoretical studies we looked at a variety of population metrics to help explain how signal filtering by the GCL improves cerebellar learning and why it ultimately fails as the GC threshold is increased.

The first set of metrics related to pattern separation: dimensionality (Dim), the number of explanatory principal components (PCs), spatiotemporal sparseness (STS), and population variability (See methods for details). (Although STS is a measure of sparseness, it represents an idealized form of separability where GCs represent unique temporal epochs that do not overlap, providing a perfect basis set when maximized, thus is grouped with pattern separation metrics). Dim, PCs, and STS showed non-monotonic relationships with threshold and peaked at thresholds ranging between 0.5 and 1.5 (Fig. 5 A, B), while population variability decreased with increasing thresholds (Fig. 5C). Intuitively, this relationship captures the effect of low thresholds allowing GC activity to relay the mean input, with no pattern separation occurring, and thus minimizing pattern separation metrics. With increasing threshold, GC activity is driven by coincidence detection leading to high dimensional population output. At high thresholds, inputs rarely summate to threshold, leading to lost representation that drives a roll-off in pattern separation within the population. Notably, Dim, PCs, and STS peaked at thresholds greater than peak learning performance, which was optimized at threshold zero, thus none of these three pattern separation metrics alone account for learning performance. Population variability (i.e. GCL variance per unit) is thought to aid classification and separability of GCL output (Cayco-Gajic et al., 2017). This metric’s decrease with increasing threshold was likely due to the decrease in overall representation by each unit due to sparsening and diminishing the dynamic range of GC rates due to threshold subtraction (Fig. 1, Fig. 5C).

**Figure 5:**
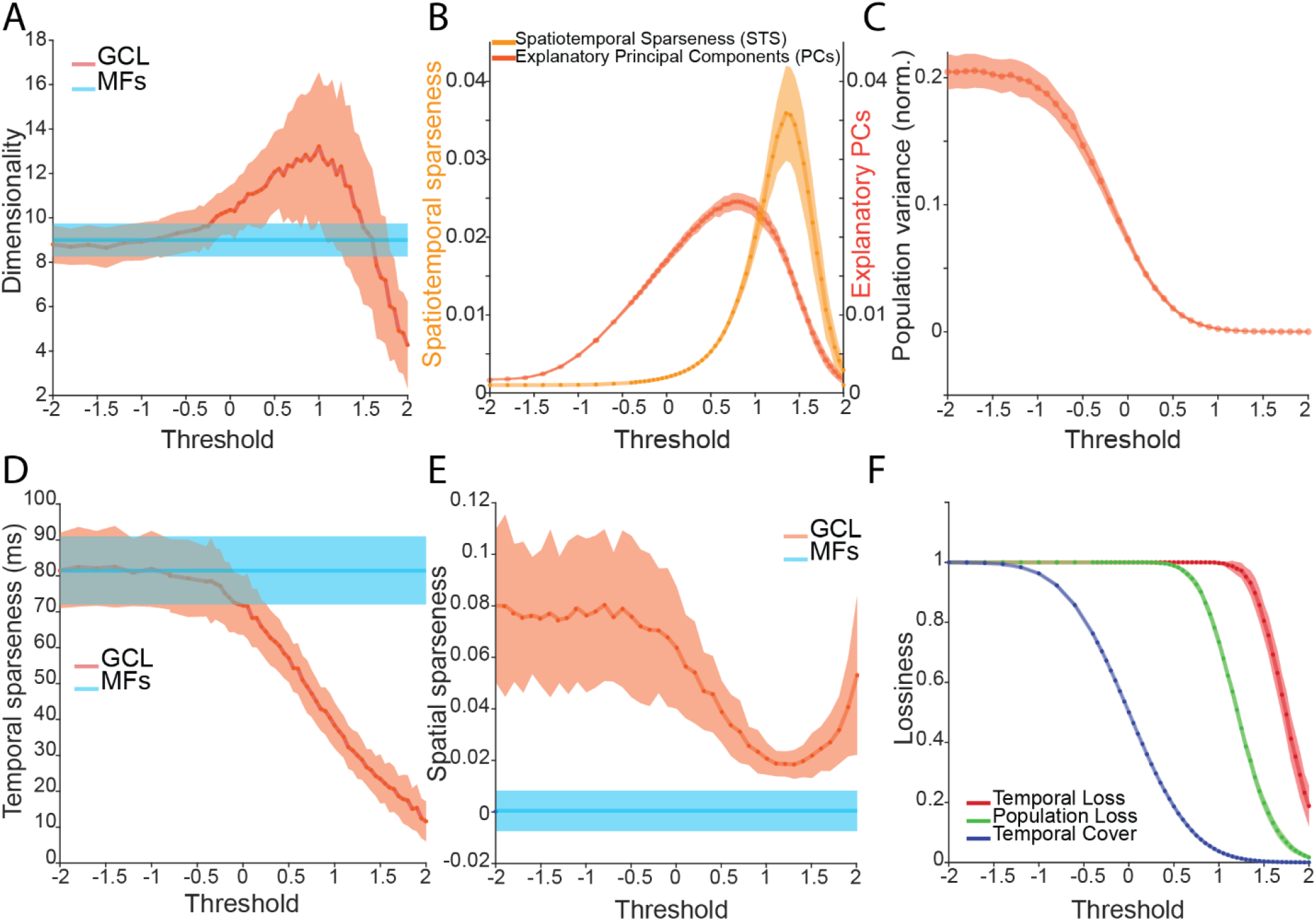
Statistical features of GCL output. **A**. GCL dimensionality (red) and MF dimensionality (blue) as a function of threshold. Note peak near a threshold of 1 for the GCL. **B**. Two metrics of pattern separation in GCL output -- STS (light orange) and PCs (dark orange) -- as a function of threshold. Note peaks near 1.5 and 0.5, respectively. **C**. The sum of GCL variance produced by the model as a function of threshold. Note monotonic decrease with threshold. **D**. Temporal sparseness as a function of thresholding. Note monotonic decrease in GCL with thresholding. **E**. Mean pairwise correlation of the population plotted as a function of threshold. Note trough near 1. **F**: Three forms of lossiness in GCL output as a function of threshold. Each metric had differential sensitivity to thresholding but note that all decrease with increasing threshold. Across metrics, function maxima and minima ranged widely and were not obviously related to thresholds of optimized learning.

The second set of metrics are related to sparse representations: temporal sparseness and spatial sparseness. Temporal sparseness – defined by the exponential decay of GC autocovariance, where smaller values typify signals that change quickly with time -- decreased as a function of threshold because of sparsened representation at higher thresholds (Fig. 5D). The mean pairwise GC correlation, (Fig. 5E) i.e. spatial sparseness, shared a drop-off after a threshold of 0, but increased again at high thresholds because only a few MF signals were retained at high threshold and thus were highly correlated. By experimental design, decorrelation was already maximized in OU inputs. Similar to the pattern separation metrics, these sparseness metrics did not show an obvious relationship to the U-shaped learning performance seen in Fig. 2A.

Finally, we examined three metrics of lossiness defined to quantify (1) the fraction of the total epoch with no activity in any GC unit (e.g. with “temporal lossiness” of 0.1, 10% of the total epoch has no activity in any GCs) (2) the proportion of granule cells with any activity over the entire epoch (“population lossiness”) (3) the mean fraction of the epoch in which each granule cell is active (“temporal cover”). Not surprisingly, each lossiness metric increased with high thresholds (Fig. 5F). However, despite diminishing activity in individual GCs with increasing threshold, (the blue curve Fig. 5F), each GC was resistant to becoming completely silent (green curve drop, Fig. 5F), owing to a few dominant inputs.

Notably, none of these metrics alone obviously tracked the U-shaped learning performance (Fig. 2A). However, collectively, these descriptive statistics of model GCL population activity set the stage for analyzing how information preprocessing by the basic GCL architecture relates to learning time series, explored below.

### Optimization of learning through GCL transformations

With the knowledge that thresholding drives changes both in learning time series (Fig. 2, 3) and GC population metrics that are theorized to modulate learning (Fig. 4, 5), we next directly investigated the relationships of these metrics to learning performance. To test this, we used a LASSO regression method to identify learning performance-driving variables taken from the metrics described in Figures 4 and 5 (Fig. 6A, C). Using the output of the LASSO model, we found that a three-term model using the most explanatory variables -- STS, the number of explanatory PCs and variance retained (Fig. 6B, C, D) -- accounted for 91% of learning variance. The three-term model performance is plotted against the observed MSE over a range of thresholds in Fig. 6D, showing strong similarity.

**Figure 6.**
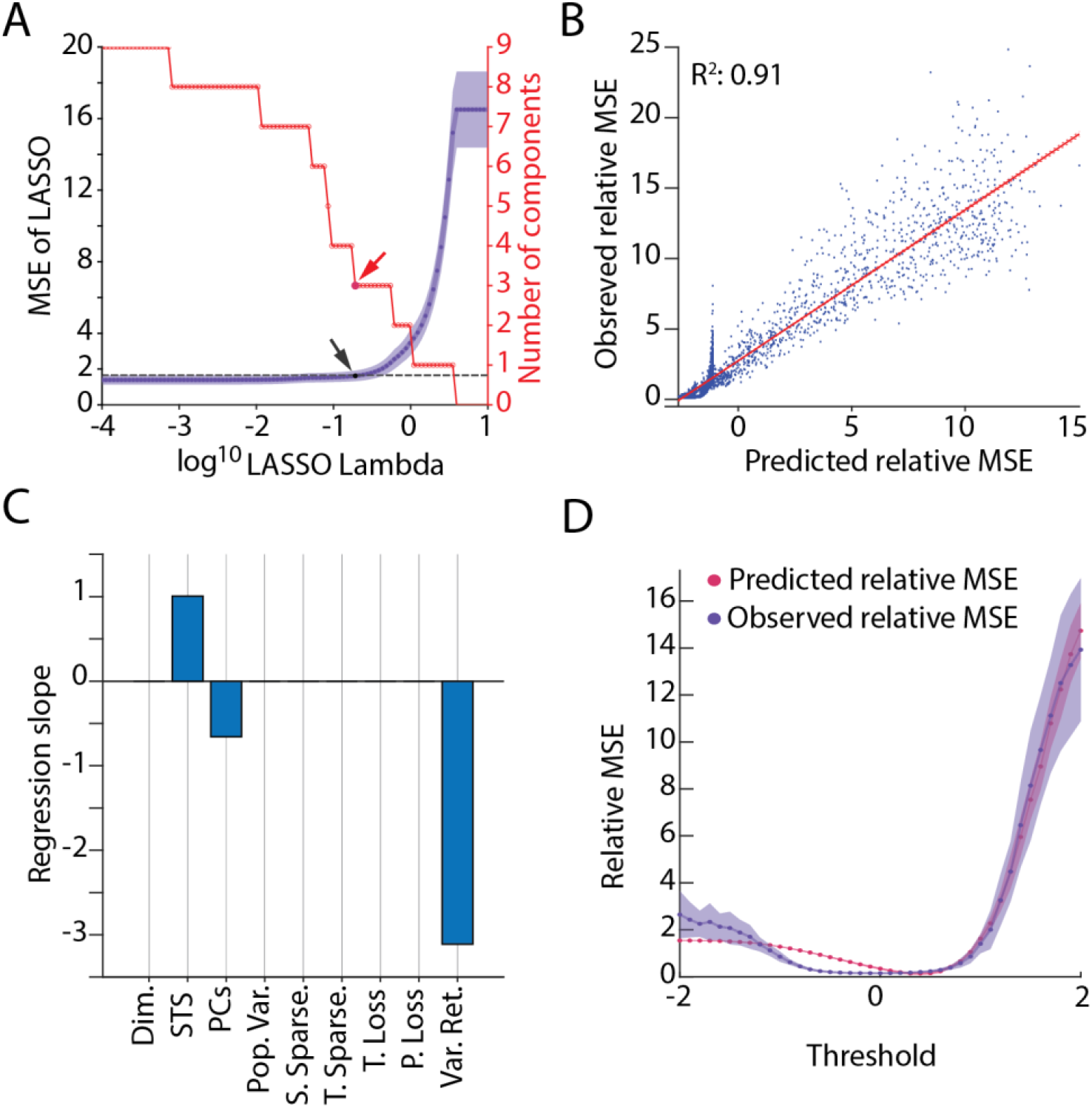
Relationship between sparseness metrics and MSE. **A**. LASSO regression model selection as a function of progression of the Lambda parameter (penalty applied to regressor selection). The removal of regressors with increasing Lambda (red steps) selected from the following potential regressors: dimensionality (Dim.), spatiotemporal sparseness (STS), explanatory principal components of the GC population (PCs), population variability (Pop. Var.), spatial sparseness (S. Sparse.), temporal sparseness (T. Sparse.), temporal lossiness (T. Loss.), population lossiness (P. Loss), and input variance retained (Var. Ret; Figure 4). Arrow shows selection point of LASSO regression MSE using “1SE” (1 standard error) method (see Methods, purple lines, black dot and arrow indicating the selected model, with red arrow showing selection point in the parameter reduction plot, red). **B**. Relationship between LASSO model (predicted relative MSE) against the observed relative MSE (ratio of GC MSE to MF alone MSE) with fit line and variance explained by regression (R^2^ = 0.91) **C**. Regression slopes of the selected LASSO model from A, showing that STS, PCs, and Input Variance Retained are the selected regressors, with Var. Ret. being the largest contributing factor. All factors normalized to a normal distribution for comparison. **D**. The output of the selected model and the observed MSE plotted against threshold for a comparison of fits, demonstrating high accuracy in the 0-2 range, but less accuracy in the -2-0 range.

These results were somewhat surprising given prior studies showing benefits of population sparseness or decorrelation to learning. To interrogate this seeming disparity, we introduced fictive GCL population activity that had specific statistical features as inputs to P-cells. Consistent with previous reports, decorrelation and temporal sparseness improved learning accuracy, with complete decorrelation and temporally sparse supporting the best performance (Fig. 6 - figure supplement 1; Cayco-Gaijic et al., 2018). Thus, on their own, population, temporal and idealized spatiotemporal sparseness do modulate learning when their contribution is independent, but these features nevertheless do not emerge as features in the naturalistic GCL model as statistical properties that drive performance of time series. This property is a consequence of temporal sparseness and decorrelation covarying with lossiness (captured by the variance retained metric), which drives down performance. Rather, the statistical features produced by the model GCL with naturalistic inputs that best explain learning are the number of explanatory PCs, STS, and the amount of input variance retained -- metrics that may align well with recently described GC population activity during locomotion (Lanore et al., 2021).

**Figure 6 figure supplement 1:**
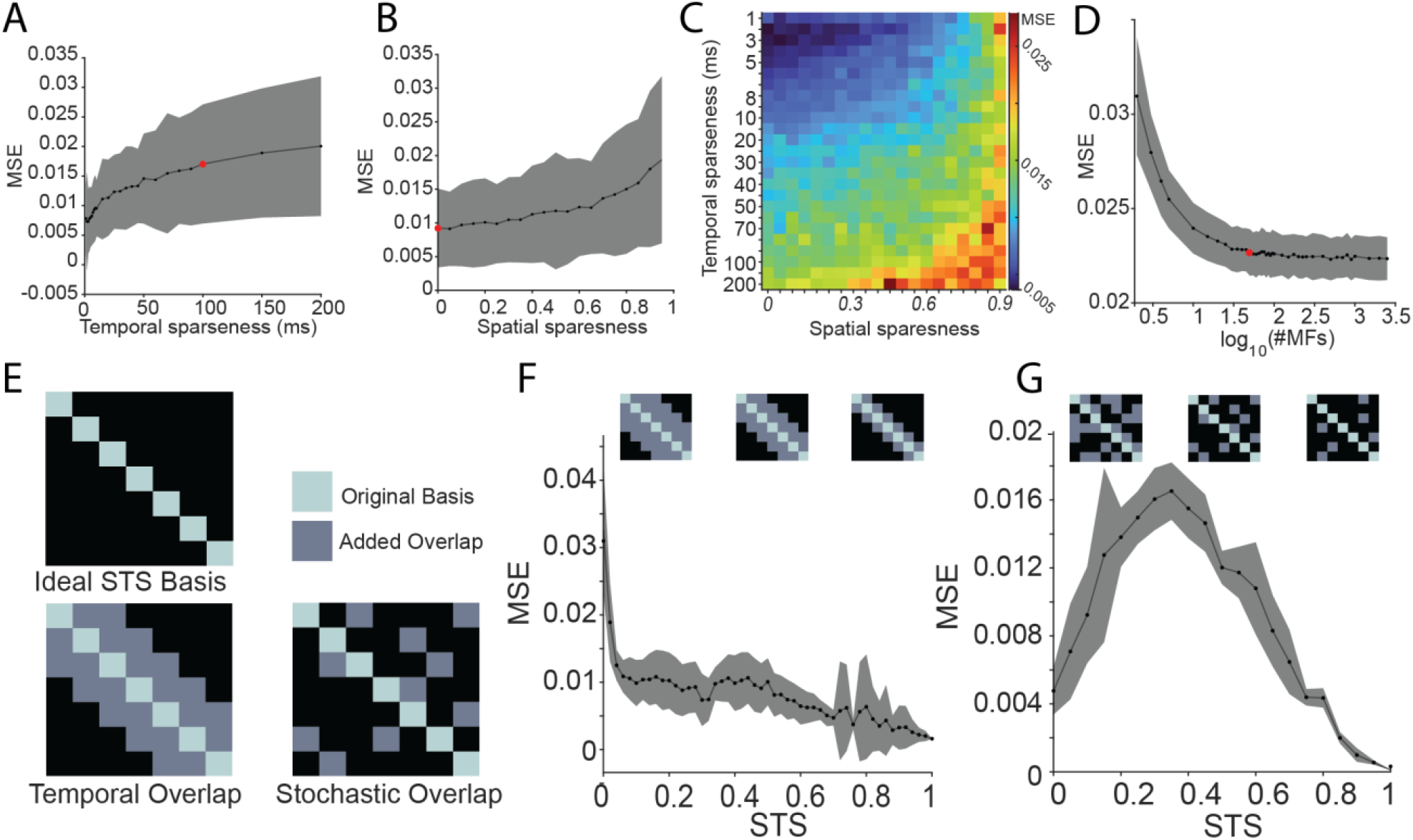
GC population statistics regulate learning accuracy when independently controlled. Fictive population activity with structured statistics were introduced to P-cells to explicitly test the roles of population decorrelation and structured spatiotemporal sparseness on learning. **A-B**: Learning performance (MSE) as a function of temporal sparseness (i.e. autocovariance tau) or spatial sparseness (i.e. population correlation). Red dots on A and B indicate values used for input model to GCL in Figs 2, 5, and 6. **C:** Matrix of effects on MSE when modulating temporal spareness via tau, and spatial sparseness via population correlation. Lower values for both (cooler colors) indicate the best learning accuracy. **D:** The results of these analyses support the idea that GCL filtering benefits learning through transformation of statistical structure fed to the P-cell. A remaining caveat was that the number of granule cells far exceeded the number mossy fibers, raising the question of whether the learning advantage conferred by the GCL is merely a consequence of this difference. To test this, we fed MFs directly to the P-cell units and varied their numbers between ranges of 2 to 3000. While learning accuracy improved with more MFs, asymptotic MSE values were lower than the GCL, indicating that the filtering properties of GCL are indeed important for this learning task. Figure plots the MSE as a function of the number of inputs to Purkinje cells, showing that too few MFs are insufficient to produce accurate learning, but having a large number makes little difference beyond 10^1.5^ ∼= 31 MFs. **E:** To test how the uniqueness of individual unit activity across time contributes to learning we selected population activity that varied in STS from a bank of simulations. Two distribution structures were tested. The first maximized granule cell uniqueness in time and temporal organization – e.g. each granule cell is active only once during the epoch and only one granule cell is active at a given time, such that the population histograms resemble a ‘staircase’ (“ideal STS basis”). Overlap of active granule cells drives decreases in computed STS, or wider steps in the staircase (“temporally linked overlap”). The second class of STS maximized uniqueness without requiring temporal organization – e.g. any slice of time is unique, but an individual granule cell can occupy an arbitrary number of time bins (“stochastic overlap”). STS drops when a given granule cell activity occupies more time bins, reducing the uniqueness of the granule cells contribution to the population. Figure shows schematic diagram of these different types of spatiotemporal sparseness, with structured overlap “temporal overlap” and “stochastic overlap” illustrating different ways populations could differ. **F:** Effect of STS on MSE, where overlap between units is always local to a particular time point, so that units are only active at a particular continuous temporal range, showing a monotonic decrease in error as STS approaches 1. **G**: Same as F, but the temporal location of overlap between units is random, showing best learning accuracy at STS = 1, and good but less accurate learning at STS = 0. When overlap was decoupled from time in the stochastic overlap case, error was reduced at both maximal and minimal STS simulations with the highest error occurring at intermediate STS values. This may be because the gradient descent algorithm is able to use dense, variable signals, like those seen in very low STS value GCL outputs, to learn essentially as well as the high STS values which have strong isolation in individual unit representation and are guaranteed to be good for learning.

### GCL properties that enhance learning in naturalistic tasks

Together, these models suggest that the GCL can reformat random inputs suitable to support rapid and accurate learning of time-series. The real cerebellum is topographically organized along multiple parasagittal output modules (Apps and Garwicz, 2005; De Zeeuw, 2020). This organization suggests segregated afferents with specific statistical structure could refine specific behaviors. To examine whether different population statistical features might support distinct learning tasks, we utilized the model to perform a series of naturalistic cerebellar tasks: vestibulo-ocular reflex (VOR) phase adaptation (Ito et al. 1974), temporal interval learning (Narain et al., 2018) and kinematic encoding (Herzfeld et al., 2015).

We speculated that the nature of these tasks might influence the contribution of components of the model to learning accuracy. For example, when VOR is kept in phase, it makes intuitive sense that retention of vestibular input, inherently in-phase with the motor output, would be valuable, with reweighting of GC representations of inputs giving rise to amplitude learning as in VOR gain adaptation. However, if the phase is offset, the relationship between vestibular input and ocular output requires complex mapping (Fig. 7A, top middle inset) and selection of GCs representing sparsened OU processes may be selected instead to allow for reconstruction from high-dimensional outputs. The GCL model supported learning of VOR at all phases, but MFs showed especially poor performance in pi/2 phase shifts (Fig. 7A, ‘out-of-phase’). As a result of this reliance on GCL reformatting, we predicted that the contribution of ‘variance retained’ to learning should decrease depending on the phase shift. In other words, the extent to which the input was inherently related to the output would be of scalable importance. We tested the relationship of input variance retention and phase offset using RIDGE regression (which preserves even small contributions of regressor variables to the model in comparison to LASSO) and found that for in-phase and anti-phase learning input variance retention accounted for most of learning, reflected in large slope coefficients, whereas input retention decreased as an important variable in out-of-phase learning, with shallow slope coefficients (Fig. 7B). Furthermore, the relative magnitude of the slope magnitude of variance retrained is reduced in out-of-phase conditions compared to in-phase and anti-phase (Fig. 7C). This suggests that the learning rule can utilize information preserved by the GCL, as in in-phase learning, but, if necessary, it can learn using information that is so highly reformatted that it no longer retains the original vestibular information.

**Figure 7.**
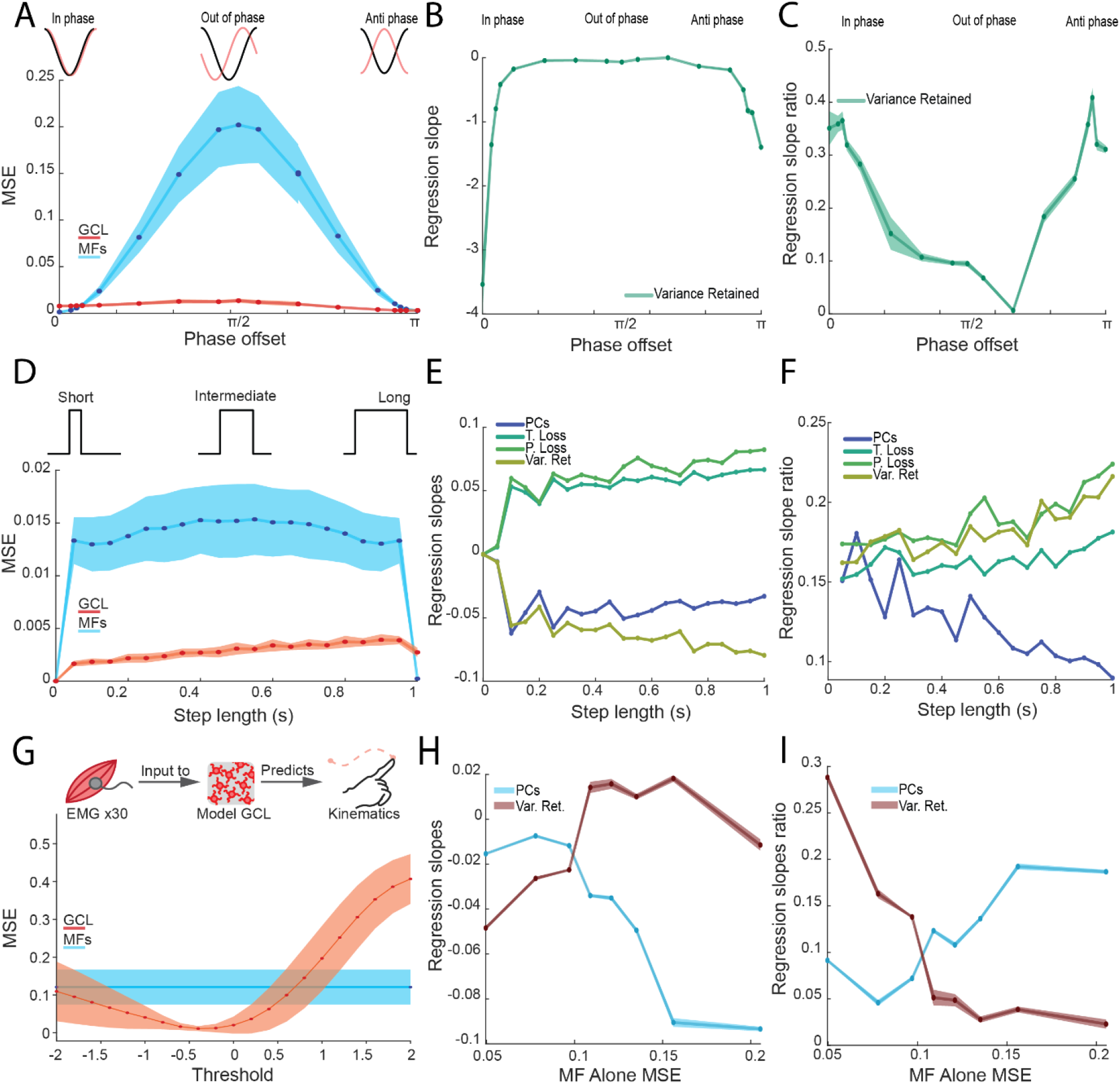
Task-dependent relationships between granule cell population statistics and learning. **A**. Task structure of a phase-offset VOR-like task (top) and learning performance as a function of phase-offset for GCL and MFs alone (bottom). Here, the phase between the input function and the target function varies between 0 and pi. GCL (red) or MFs alone (blue) were used as inputs to learn the task. As the difference in phase between inputs and targets approaches pi/2 (out of phase), performance from MF alone degrades while GCL performance remains accurate and stable. **B**. RIDGE regression slopes of the input variance retained (Var. Ret.) metric as a function of phase offset. Variance retained slope is large when phase offset is in the ‘in phase’ and ‘anti-phase’ regions of the task, but is otherwise minimized, suggesting that the utility of this statistical feature varies depending on task. **C**. Same data as B but normalized to show the relative proportion of all slope magnitudes accounted for by Var. Ret. (slope magnitude of Var. Ret divided by the sum of all slope magnitudes). Var. Ret. is a primary regressor for ‘in phase’ or ‘anti-phase’ learning. **D**. Task structure (top) and learning performance (bottom) of an interval estimation task, where the model is tasked with learning a step function that varies in length. GCs (red) and MFs alone (blue) were used as inputs to the P-cell. As the interval lengthens, learning using MFs alone was generally poorer than using the GCL. **E**. RIDGE regression slopes of 4 variables (Var. Ret., T. Loss, P. Loss, PCs) as a function of step length, showing that slopes of lossiness-related metrics (P. & T. Loss, and Var. Ret.) increase in magnitude as the step length increases, whereas slope magnitude of PCs decreases. **F**. Same as E but showing the relative proportion of all slopes accounted for by these 4 regressors. **G**. Schematic of underlying dataset using recorded EMG as an input to the model GCL to predict kinematics (top). Learning performance of model using EMG alone (MFs; blue) or GCL (red) across varying thresholds. The GCL outperforms MFs alone at a threshold range similar to that observed in Fig. 2. **H**. RIDGE regression slopes of Var. Ret and PCs metrics as a function of the learning performance achieved by using MFs (i.e. EMG) alone, showing that Var. Ret. is a stronger driver of performance when MFs alone supported accurate learning, but not when MFs alone supported poor learning (higher MSE). PCs show the opposite trend, increasing in slope magnitude when MFs alone supported poor learning. **I**. Same as H, but showing the relative proportion of slope magnitude accounted for by Var. Ret. and PCs.

Cerebellar timing tasks, such as delay eyeblink conditioning, involve time estimation over intervals spanning 100-500 *ms*. Models of delay eyelid conditioning suggested that the cerebellum represents the time interval through decomposition of an invariant signal into many signals that tile across time. This hypothesis provides an interesting test of whether lossiness differentially affects behavioral outcomes depending on whether short or long intervals are being estimated, defined here as the duration of an ‘on’ signal. If we assume that an ideal temporal basis (akin to the ‘staircase’ representation in Fig 6 - figure supplement 1E) represents different points in time of the stimulus, one might speculate that lossiness in populations representing long intervals would be more detrimental than in populations representing short intervals -- given that only the temporally aligned subsection of the input is relevant to the output response and the rest is discarded or ignored. We tested this prediction by systematically altering the length of a step target function to occupy 0% to 100% of the response epoch using OU processes as inputs. The model using a GCL was able to perform this task more accurately than with MF inputs alone (Fig 7D), and the magnitude of slope for lossiness-related metrics increased with interval duration (Fig. 7E), suggesting that learning short intervals is less sensitive to lossiness than learning long intervals. The relative contribution of lossiness metrics to the overall regression performance also increased with step duration compared to PCs (Fig. 7F), suggesting that lossiness-related metrics have a more powerful influence on learning outcomes as a function of increasing duration that is not true of pattern separation metrics like PCs.

We next asked whether naturalistic input statistics, derived from electromyogram (EMG) signaling, could support learning. We used EMG signals from human subjects in a point-to-point reaching task as MF inputs, and tested whether the model could learn associated limb kinematics from this input (Fig. 7G; Delis et al. 2018; Tseng et al. 2007; Miall and Wolpert 1996; Wolpert et al., 1998). The GCL was able to produce more accurate predictions of the kinematics when compared to the EMG as MF inputs alone, and the range of thresholds which produced the best accuracy were comparable to the previous findings (Fig. 3A), but were slightly negatively shifted, suggesting retained variance of inputs might be beneficial to learning kinematics from associated muscle activity.

Finally, since EMGs used as MF inputs to the model had some level of baseline utility in predicting kinematics based on their intrinsic relationships, (reflected in MFs alone MSE varying between 0.04 and 0.22), we next asked whether this influenced which features of the GCL output were most related to learning. In keeping with intuition, when MF based learning was excellent (low MSE), the slope of the variance retained metric was highest (Fig. 7H, I, blue). Conversely, when MF based learning was poor (high MSE) variance retained slopes dropped. Interestingly, a few GCL population metrics became more important for learning as MF MSE worsened, such as the number of explanatory PCs (Fig. 7H, I, maroon). Together this suggests that different pattern separation features of GCL reformatting may serve learning under different conditions, with Purkinje cells using diverse ‘pattern separation’ features depending on the task and input statistics. When intrinsic relationships are valuable, variance retained is an important population statistical feature; when they are more arbitrary, pattern separation features are more valuable for learning relationships between the inputs and output. This shifting landscape was a general feature of our models (Fig. 6 & 7), suggesting that “pattern separation” by the GCL is not one universal transform that has broad utility. This observation raises the possibility that regional circuit specializations within the cerebellar cortex, such as density of unipolar brush cells (Dino et al. 2000), Golgi cells, or neuromodulators could bias GCL information reformatting to be more suitable for learning of different tasks.

## Discussion

Here we asked a simple question: how does the cerebellar granule layer support temporal learning? This question has captivated theorists for decades, leading to a hypothesis of cerebellar learning that posits that the GCL reformats information to best suit associative learning in Purkinje cells. Recent work has called many of these foundational ideas into question, however, including whether GCL activity is sparse; high dimensional; and what properties of ‘pattern separation’ best support learning (Wagner et al., 2017; Giovannucci et al., 2017; Knogler et al., 2017; Cayco-Gajic et al., 2017; Gilmer and Person 2017). To reconcile empirical observations with theory, we hypothesized that input statistics and task structures influence how the GCL supports learning. Here, we used naturalistic time-varying inputs to a model GCL and identified pattern separation features that supported learning a time series prediction task, with an arbitrary but temporally linked input-output mapping, recapitulating important features of physiological cerebellar learning tasks. This formulation eliminates the possibility of trivial dimensionality changes improving classification performance, thus approaching naturalistic challenges faced by the real circuit. Several important observations stemmed from these simulations: (1) with naturalistic input statistics, the GCL produces temporal basis sets akin to those hypothesized to support learned timing with minimal assumptions; (2) this reformatting is highly beneficial to learning; (3) maximal pattern separation does not support the best learning; (4) rather, tradeoffs between loss of information and reformatting favored best learning at intermediate network thresholds; and finally (5) different “cerebellar” tasks utilized different GCL population statistical features to optimize performance. Together these findings provide insight into the granule cell layer as performing pattern separation of inputs that transform information valuable for gradient descent-like algorithms (akin to Purkinje cell learning rules), but with idiosyncratic population statistics supporting different tasks. This observation makes predictions about the regional specifications that occur across the layer that may specially subserve diverse behaviors.

### Emergence of spatiotemporal representation and contribution to learning

A perennial question in cerebellar physiology is how the granule cell layer produces temporally varied outputs that could support learned timing (Mauk and Buonomano 2004). While cellular and synaptic properties have been shown to contribute (Chabrol et al., 2015; Duguid et al, 2012; Guo et al., 2021; Crowley et al., 2009; Rudolph et al., 2015; Buonomano and Mauk 1994; Kanichay and Silver 2008; Simat et al., 2007; Mapelli et al., 2009; Rossi et al., 1996; Gall et al., 2005; Armano et al., 2000; Rizwan et al. 2016; Tabuchi et al., 2019; D’Angelo and De Zeeuw 2009), we observed that with naturalistic inputs, temporal basis set formation is a robust emergent property of the threshold-linear input-output function of granule cells receiving multiple independent time-varying inputs (Fig. 1B). But is this reformatting beneficial to learning? We addressed this question by comparing learning of a complex time-series in model Purkinje cells receiving either mossy fibers alone or reformatted output from the GCL. We found that indeed the GCL outperformed MFs alone in all tasks (Figs. 2, 3, 7). Nevertheless, we wondered what features of the population activity accounted for this improved learning. While sparseness, decorrelation, dimensionality and lossless encoding have been put forward as preprocessing steps supporting learning, we found that none of these alone accounted for the goodness of model performance. Rather, disparate pattern separation metrics appear to strike a balance between maximizing sparsenesses without trespassing into lossy encoding space that severely, and necessarily, degrades learning of time-series.

Moreover, the value of different population metrics to learning varied with the specific task -- with some tasks relying on input retention for best performance, others relying on absence of lossiness, and some requiring pattern separation to accomplish accurate predictions (Fig. 7). For instance, when input statistics are well suited to learning the specific task, as in in-phase VOR, preservation of input variance drives best performance (Fig. 7A-C). Importantly, although the properties of the GCL selected to improve learning varied across tasks, the underlying architecture of the GCL and thresholding did not. This suggests that the output of the GCL is well structured to support a variety of tasks. Thus, Purkinje cells are able to make use of a spectrum of information formats that, depending on task requirements, are selected to serve best learning.

These observations are interesting in light of a long history of work on granule layer function. Marr, Albus, and others proposed that the granule cell layer performs pattern separation useful for classification tasks. In this framework, sparseness is the key driver of performance, and could account for the vast number of granule cells. Nevertheless, large-scale GCL recordings unexpectedly showed high levels of correlation and relatively non-sparse activity (Wagner et al., 2017; Giovannucci et al., 2017; Knogler et al., 2017). Despite methodological caveats, alternate recording methods seem to support the general conclusion that sparseness is not as high as originally thought (Lanore et al. 2021; Kita et al., 2021; Gurgani and Silver 2021). Indeed, subsequent theoretical work showed that sparseness has deleterious properties (Cayco-Gajic et al., 2017; Billings et al., 2014), also observed in the present study, that may explain dense firing patterns seen *in vivo*. Here we found that the best learning occurred when individual granule cell activity occupied around half of the observed epoch (Fig. 5F, blue trace), achieved with intermediate thresholding levels. We also observed temporal organization that is consistent with the firing patterns observed *in vivo*. While these findings seem to suggest that sparseness is not the ‘goal’ of GCL processing, our findings and others (Litwin-Kumar et al., 2016; Cayco-Gajic et al., 2017) suggest that pattern separation broadly is a positive modulator of GCL support of learning processes.

Previous work (Sanger et al., 2020) proposed that time-series prediction was possible with access to a diverse set of geometric functions represented in the GC population. However, that study left open the question of how such a diverse collection of basis functions would emerge. The GCL model used here minimized free parameters by incorporating very few independent circuit elements, suggesting that a single transform is sufficient to produce a basis set which is universally able to learn arbitrary target functions. We used a simple threshold-linear filter with a singular global threshold that relied on sparse-sampling to produce spatiotemporally varied population outputs. This simple function worked to support learning at a broad range of inputs and thresholding values, ultimately allowing the Purkinje cells downstream to associate the spatiotemporally sparser inputs with feedback to learn arbitrary, and often quite difficult, target functions. The emergence of this basis set is remarkable given the very simple assumptions applied, but is also physiologically realistic, given the simple and well characterized anatomical properties of the MF divergence and convergence patterns onto GCs, which are among the simplest neurons in the brain (Jakab and Hamori, 1988; Palay and Chan-Palay, 1974; Palkovits et al., 1971). Although we suggest that the key regulator of thresholding in the system is the feedforward inhibition from Golgi cells, many factors may regulate the transformation between input and GC output in the network, allowing for multiple levels and degrees of control over the tuning of the filter or real mechanism that controls the outcomes of GCL transformations. Golgi cell dynamics may prove critical for enforcing the balance between pattern separation metrics and lossy encoding (Hull 2020) thus are critical players in mean thresholding found here to optimize learning. Additional mechanistic considerations may also play a role, including short-term synaptic plasticity (Chabrol et al. 2015) and network recurrence (Gao et al. 2016; Houck and Person 2014; 2015; Judd et al., 2021), allowing for a more nuanced and dynamic regulatory system than the one shown here.

### Recapturing input information in the filtered GCL output

Two schools of thought surround what information is relayed to Purkinje cells through GCs. Work in the oculomotor cerebellum and flocculus suggests that Purkinje cells inherit virtually untransformed information encoding eye velocity and visual motion, integrated in P-cells as positional signals (Herzfeld et al., 2020; Krauzlis and Lisberger, 1991). Alternatively, the implication of theories of Marr and Albus suggest that input information is so sparsened that Purkinje cells receive only a small remnant of the sensorimotor information sent to the cerebellum. These divergent views have never been reconciled to our knowledge. We addressed this disconnect by determining the fraction of MF input variance recoverable in GCL output. Interestingly, the GCL population retains sufficient information to recover more than 90% the input variance despite filtering out 50% or more of the original signal (Fig. 4). This information recovery is achieved at the population level and thus requires sufficient numbers of granule cells so that the subset of signals that are subthreshold are also super-threshold in other subsets of GCs through probabilistic integration with other active inputs. While variance recovery is not a true measure of mutual information, it is indicative of the utility that the intersectional filtering performed by the GCL. The expansion of representations in the GCL population achieved by capturing the coincidence of features in the input population creates a flexible representation in the GCL output that has many beneficial properties, including the preservation of information through some degree of preserved mutual information between the GCL and its inputs.

### Enhanced learning speed

Our model not only improved learning accuracy, but also speed, compared to MFs alone (Fig. 3). Both learning speed and accuracy progressed in tandem: threshold parameter ranges that enhanced overall learning speed also minimized mean squared error, suggesting that speed and accuracy are enhanced by similar features in GCL output. Learning speed was well described by a double exponential function with a slow and fast component. This dual time course in the model with only one learning rule is interesting in light of observations of behavioral adaptation that also follow dual time courses (Herzfeld et al., 2014; Smith et al., 2006). Some behavioral studies have postulated that these time courses suggest multiple underlying learning processes (Yang and Lisberger, 2014). Our model indicates that even with a single learning rule and site of plasticity, multiple time-courses can emerge, presumably because when error becomes low, update rates also slow down.

Another observation stemming from simulations studying learning speed was that the behavior of the model varied as a function of the learning ‘step size’ parameter of the gradient descent method (Fig 3 – Fig. Supplement 1). The step size -- ie. the, typically small, scalar regulating change in the weights between GCs and P-cells following an error -- determined the likelihood of catastrophically poor learning: when the step size was too large, it led to extremely poor learning because the total output ‘explodes’ and fails to converge on a stable output. Nevertheless, the model tolerated large steps and faster learning under some conditions, since the threshold also influenced the likelihood of catastrophic learning. Generally, higher thresholds prevented large weight changes from exploding, suggesting that sparse outputs may have an additional role in speeding learning by supporting larger weight changes in Purkinje cells. Indeed, appreciable changes in simple spike rates occur on a trial-by-trial basis, gated by the theorized update signals that Purkinje cells receive, climbing fiber mediated complex spikes. These plastic changes in rate could reflect large weight updates associated with error. Moreover, graded complex spike amplitudes that alter the size of trial-over-trial simple spike rate changes suggest that update sizes are not fixed (Najafi et al., 2014; Herzfeld et al., 2020; Medina and Raymond 2018). It is possible that the amplitude of synaptic weight changes following a complex spike might be set by tunable circuitry in the molecular layer to optimize learning speed relative to the statistics of the GCL output.

Together, this study advances our understanding of how the GCL may diversify or isolate components of inputs. A number of behavioral observations might be informed by the present findings. The timecourse of learning for instance varies widely across tasks. Eyeblink conditioning paradigms require hundreds of trials to learn (Millenson 1997; Khilkevich et al., 2016; Lincoln et al., 1982), while saccade adaptation and visuomotor adaptation of reaches, which are also mediated by the cerebellum (Raymond and Lisberger; Martin et al. 1996), requires just tens of trials (Tseng et al., 2007; Shadmehr and Mussa-Ivaldi 1994; Ruttle et al., 2021; Calame et al., 2021). This discrepancy in learning rates raised the possibility that the learning algorithm used by the cerebellum is better engaged during naturalistic movements compared to time-invariant cues, such as a conditioning stimulus. Such purely time-invariant cues would be difficult, if not impossible, for our model GCL to reformat and sparsen, as they are incompatible with thresholding-based filtering of input signals used here. Supportive of this view, recent work showed that EBC learning was faster if the animal is locomoting during training (Albergaria et al., 2018). We hypothesize that naturalistic time-variant signals associated with ongoing movements inputted to the cerebellum through MFs support robust temporal pattern separation in the GCL, enhancing learning accuracy and speed, while time invariant associative signals used in typical classical conditioning paradigms result in an impoverished ‘basis’, making learning more difficult. That this feature is so robust could explain why tasks like eyeblink conditioning are so difficult to learn, sensorimotor tasks can be adapted rapidly. We speculate that the cerebellum is structured to support fast learning in situations where there are physiologically structured inputs, typified by convergent, temporally varying self-generated efference and reafference, within rich sensory and motor environments, as in normal movements during daily life.

## Methods

### Model construction

The model presented here incorporated major features of the granule cell layer (GCL) circuit anatomical organization and physiology. The features chosen for the model were the sparse sampling of inputs (GCs have just 4 synaptic input branches in their segregated dendrite complexes on average), which was reflected in the connectivity matrix between the input pool and the GCs, where each GC received 4 inputs with weights of 1/4^th^ (i.e. 1 divided by the number of inputs; 1/M) of the original input strength, summing to a total weight of 1 across all inputs. The other features were thresholding, representing inhibition from local inhibitory Golgi neurons and intrinsic excitability of the GCs. The degree of inhibition and intrinsic excitability (threshold) was a free parameter of the model, and the dynamics were normalized to the z-score of the summated inputs. This feature reflects the monitoring of inputs by Golgi cells while maintaining simplicity in their mean output to GCs. While this model simplifies many aspects of previous models of the GCL, it recreated many of the important features of those models, suggesting that the sparse sampling and firing are the main components dictating GCL functionality.

The model, in total, uses the following formulas to determine GC output:

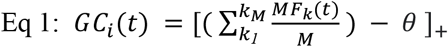

where k is a random selection of M MFs from the MF population. The inputs are summed and divided by the total number of MF inputs to the GC, M, so that their total weight is equal to 1. Unless noted as a variable, we used M = 4, reflecting the mean connectivity between MFs and GCs, and the optimal ratio for expansion recoding (Litwin-Kumar et al. 2017), and the point of best input variance retention (Fig. 4). This function is then linearly rectified, i.e. [*x*]_+_ = *x* if *x* > 0 and 0 otherwise so that there are no negative rates present in the GC activity. The *θ* function which determines the threshold mimics intrinsic excitability and feedforward inhibition was formulated as:

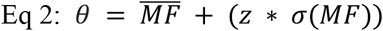

Here, a function of the mean and standard deviation of the entire MF population, z is a free parameter in the model representing the number of standard deviations from the mean, setting the minimum value below which granule cell activity is suppressed, which is the threshold value reported within this study as ‘threshold’. Note that the summated MF inputs are divided by the number of inputs per GC (N) in Eq. 1 such that their received activity relative to *θ* is proportional to the input size, M.

### Input construction

To provide a range of inputs with physiological-like temporal properties that could be parameterized, we used a class of randomly generated signals called Ornstein-Uhlenbeck Processes (OU), defined by the following formula:

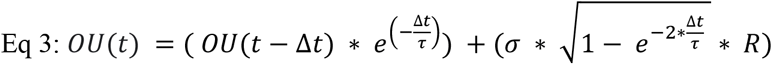

Here t is the time point being calculated, Δt is the time interval (the time base is in *ms* and Δt is 1 *ms*). *σ* is the predetermined standard deviation of the signal, and R is a vector of normally distributed random numbers. This process balances a decay term, the exponential with e raised to - Δt/*τ*, and an additive term which introduces random fluctuations. Without the additive term, this function decays to zero as time progresses. After the complete function has been calculated, the desired mean is added to the timeseries to set the mean to a predetermined value.

The vector R can also be drawn from a matrix of correlated numbers, as was the case in Fig. 6 – figure supplement 1 B & C. These numbers were produced with the MATLAB functions randn() for normal random numbers, and mvnrnd() for matrices with a predetermined covariance matrix supplied to the function. The covariance matrix used for these experiments was always a 1-diagonal with a constant, predetermined, covariance value on the off-diagonal coordinates.

### Learning accuracy and speed assay

In order to understand how the GCL contributed to learning, we constructed an artificial Purkinje cell (P-cell) layer. The P-cell unit learned to predict a target function through a gradient descent mechanism, such that the change in weight for each step was:

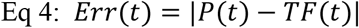

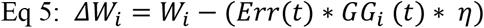

Where P(t) is the output of the P-cell at time t, TF(t) is the target function at time t, W_i_ is the weight between the Purkinje cell and the i^th^ GC, and η is a small scalar termed the ‘step size’. η was 1E-3 for GCs, and 1E-5 for MF alone in simulations shown in this study where the step size was held fixed, which was chosen to maximize learning accuracy and stability of learning for both populations. The learning process in Eq. 4 and 5 was repeated for T trials at every time point in the desired signal. The number of trials was chosen so that learning reached asymptotic change across subsequent trials. Typically, 1000 trials were more than sufficient to reach asymptote, so that value was used for the experiments in this study.

The overall accuracy of this process was determined by calculating the mean squared error between the predicted and desired function:

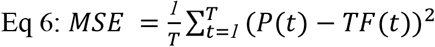

The learning speed was determined by fitting an exponential decay function to the MSE across every trial and taking the tau of the decay (See methods: Model output metrics, Time decay).

### Model output metrics

To assay the properties of the GCL output that influence learning, we measured the features of GCL output across a spectrum of metrics that have theoretically been associated with GCL functions like pattern separation or expansion, as well as optimization or cost-related metrics developed for this paper. These included: dimensionality, spatiotemporal sparseness, contributing principal components, spatial sparseness (mean population pairwise correlation), temporal sparseness (mean unit autocovariance exponential decay), population variance, temporal lossiness, population lossiness, and temporal cover.

We considered three forms of lossiness here, two related to the dimensions of sparseness considered above, time and space, and one that is a measure of sparseness on the individual GC level. Temporal lossiness is a measure of the percentage of time points that are not encoded by any members of the GCL population, essentially removing the ability of P-cells to learn at that time point and producing no output at that time in the final estimation of the target function. Increases in the value are guaranteed to degrade prediction accuracy for any target function that does not already contain a zero value at the lossy time point.

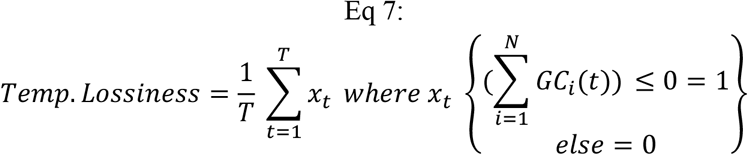

Here, T is the total number of points in the encoding epoch, the bracketed portion of the formula is a summation of inputs from all GCs (N = population size) at that timepoint. When all GCs are silent, the sum is 0, and the temporal lossiness is calculated as 1, and when all time points are covered by at least one GC, total temporal lossiness is 0.

Spatial lossiness, or population lossiness, is the proportion of GCs in the population that are silent for the entirety of the measured epoch. This is thought to reduce total encoding space and deprive downstream P-cells of potential information channels and could potentially impact learning efficacy. It is defined as:

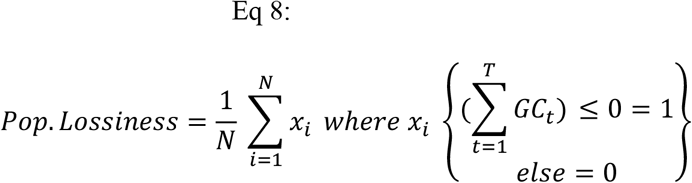

Here, N is the total population size of the GCL, and the bracketed portion of the formula is a sum of the activity of GCs across all timepoints, such that if a GC is silent across all timepoints x_i_ is calculated as 1, indicating the ‘loss’ of that GC unit’s contribution. When all GCs are silent, population lossiness is 1, and when all GCs are active for at least one time point, population lossiness is 0.

Additionally, we looked at the mean sparseness of activity across the population by measuring the ‘coverage’ or proportion of time points each GC was active during, defined as:

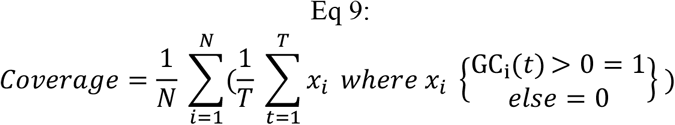

As before, N is the number of cells in the population and T is the total length of the epoch. The bracketed function counts the number of time points where GC_i_ is active, and divides that by the total time period length to get the proportion of time active. This value is summed across all GCs and divided by N to calculate the average coverage in the population. This value has strong synonymy with population variance, so it was not used for fitting assays in later experiments (Fig. 6), but reflects the effect of thresholding on average activity in the GCL population.

Dimensionality is a measure of the number of independent dimensions needed to describe a set of signals, similar in concept to the principal components of a set of signals. This measure is primarily influenced by covariance between signals, and when dimensionality approaches the number of signals included in the calculation (n), the signals become progressively independent. The GCL has previously been shown to enhance the dimensionality of input sets and does so in the model presented here too. Dimensionality is calculated with:

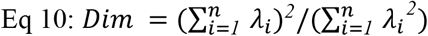

Provided by Litwin-Kumar, et al, 2016. This is the ratio of the squared sum of the eigenvalues to the sum of the squared eigenvalues of the covariance matrix of the signals.

Spatiotemporal Sparseness (STS) was a calculated cost function meant to measure the divergence of GC population encoding from a ‘perfect’ diagonal function where each GC represents one point in time and does not overlap in representation with other units. This form of representation is guaranteed to produce perfect learning, and transformations between the diagonal and any target function can be achieved in a single learning step, making this form of representation an intriguing form of GCL representation, if it is indeed feasible. We calculated the cost as:

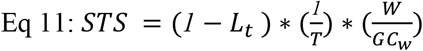

Where (1 –L_t_) is the cost of temporal lossiness, defined above (Eq. 7), and T is the total length of the epoch. W is the number of unique combinations (termed ‘words’, akin to a barcode of activity across the population), of GCs across the epoch at each point of discrete time, and GC_w_ is the average number of words each GC is active at all within the time-bins chosen (e.g. a binary representation of GC activity). The intuition used here is that when there is no temporal lossiness, all points in time are represented, leading the 1 –L_t_ term to have no effect on the STS equation, and when W, the number of unique combinations of GC activities is equal to T, then each point in time has a unique ‘word’ associated with it. Finally, when GC_w_ is 1, W/GC_w_ is equal to W, which only occurs when each GC contributes to a single word. When these conditions are met, STS = 1, otherwise when GCs contribute to more than one word, GC_w_ increases and W is divided by a number larger than 1, decreasing STS. Alternately, when there are not many unique combinations, such as when every GC has the exact same output, W/GC_w_ is equal to (1/T), decreasing STS. Finally, because lossiness causes the occurrence of a ‘special’, but non-associable, word, we multiplied the above calculations by (1 –L_t_) to account for the effect of the unique non-encoding word (i.e. all GCs inactive) on distance from the ideal diagonal matrix.

Mean temporal decay, i.e. temporal sparseness, is a measure of variance across time for individual signals, where a low value would indicate that the signals coherence across time is weak, meaning that the signal varies quickly, whereas a high value would mean that trends in the signal persist for long periods of time. This value is extracted by fitting an exponential decay function to the autocovariance of each unit’s signal and measuring the tau of decay in the function:

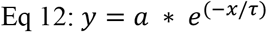

This is converted to the ms form by taking the ratio of 1000/τ. *y* here τ is a description of the autocovariance of the activity of a MF or GC signal, so when the descriptor *τ* is a large number, the decay in autocovariance is longer, or slower, when *τ* is a small number, the autocovariance across time decays more quickly, making the change in activity faster.

While dimensionality and STS are metrics rooted in a principled understanding of potentially desirable properties of population encoding, the gradient descent algorithm can extract utility from population statistics that are much noisier and correlated than the ideal populations that dimensionality and STS account for. To measure a more general pattern separation feature in GCL output that could still be associated with the complex target function, we turned to principal component analysis (PCA) with the intuition that components which explain variance in the GCL output could be utilized by the downstream Purkinje cell units to extract useful features from the input they receive (Lanore et al., 2021). We parameterized the utility of this measure by taking the proportion of the PCs derived from the GCL output which explained variance (of the GCL output) in that population by more than or equal to 1/N, where N is the number of GCs, suggesting that they explain more variance than would be expected from chance.

Population correlation, was measured by taking the mean correlation between all pairwise combinations of GCs using the corr() function in MATLAB and excluding the diagonal and top half of the resultant matrix.

Population aggregate variance is a measure related to the expansion or collapse of total space covered by the encoding done by a population, and higher or expanded values in this metric are thought to assist in pattern separation and classification learning.

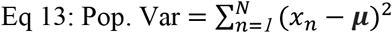

As shown in Cayco-Gajic et al. (2017). Here x is the activity of one of n cells across a measured epoch, and μ is the mean of that activity. This value is reported relative to the number of GC units, such that Pop. Var reported in Fig. 5 is normalized to Pop. Var / N.

### Variance retained assay

To test the recovery of inputs by a feedforward network with a granule cell layer (GCL), we used explained variance, *R*^2^, to quantify the quality of recovery of a sequence of normal random variables (Fig. 2) across *N*_*w*_ = 1000 numerical experiments. To distinguish this metric from the MSE and R^2^ metrics to evaluate other models in the study, we rename this ‘variance retained’. Within each numerical experiment *i*, at each time point, a vector of inputs ***x***_***t***_ of length *M* (representing the mossy fiber, MF, inputs) was drawn from an *M*-dimensional normal distribution with no correlations, ***x***_***t***_ ∼ **𝒩(0**,***I***_***M***_). This vector is then left-multiplied by a random binary matrix *W* with *N* rows and *M* columns with *n* 1’s per row and the rest zeros, followed by a threshold linearization to obtain the GCL output, ***y***_***t***_ = [*W****x***_***t***_ − ***z***]_+_ with threshold. This process is then repeated *T* = 1000 times and a downstream linear readout was fit to optimally recover ***x***_***t***_ from ***y***_***t***_. It can be shown multivariate linear regression (MATLAB’s regress() function, employing least squares to minimize mean squared error) solves this problem, identifying for each MF input stream 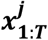, the optimal weighting *B*_*1:T*_ from the GCL to estimate 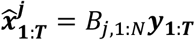. Across time *t* = 1: *T*, we then computed the squared error across the vector, 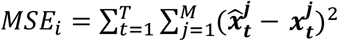 as well as the summed variance of the actual input, 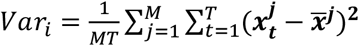, where 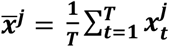 is the mean of the *j*th MF input stream. Lastly, to compute variance explained, we take 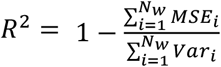, so the higher the relative mean squared error is, the lower the variance explained will be. To generate the panels in Fig. 4, we always kept the number of timepoints and experiments the same, but varied (Fig. 4B) the threshold along the axis and the number of inputs *n* per GC output; (Fig. 4C) the total number of GC outputs *N* and input per output *n*; (Fig. 4D) number of inputs *M* and outputs *N*; and finally (Fig. 4E) the number of inputs per GC output *n* along with the total number of outputs *N*.

### Independent measures generation

To determine if the sparseness measures had inherent benefits for learning, we supplemented the GCL output with OU processes with known temporal and correlational properties to examine their effect on learning accuracy (Figure 6 figure supplement 1). We varied the temporal properties by systematically varying the tau value in the exponential decay function. To vary population correlation, the random draw function in the OU process was replaced with a MATLAB function, mvnrnd(), which allowed for preset covariance values to direct the overall covariance between random samples. We used a square matrix with 1s on the diagonal and the desired covariance on all off-diagonal locations for this process and varied the covariance to alter the correlation between signals. The OU outputs from this controlled process were then fed into model P cells with randomized OU targets, as per the normal learning condition described above. To vary the effect of the input population size, the size of the supplemented population varied from 10 to 3000 using a tau of 10 and drawing from normal random numbers.

To measure the effects of STS on learning, a diagonal matrix was used at the input to a Purkinje unit, which represented population activity with an STS of 1 (see Eq 11 in Model output metrics). To degrade the STS metric, additional overlapping activity was injected either by expanding temporal representation or at random, for example, adding an additional point of activity causes inherent overlap in the diagonal matrix, increasing the GC_w_ denominator of Eq 11 to (1 + 2/N) because the overlapping and overlapped units now each contribute to 1 additional neural word. This process was varied by increasing the amount of overlap to sample STS from 0 to 1.

### GCL output metrics fits to learning

To estimate the properties of GCL output that contribute to enhanced learning of time series, we used multiple linear regression to find the fit between measures of GCL population activity and observed MSE in learning. Because there are large inherent correlations between the metrics used (dimensionality, spatiotemporal sparseness, explanatory principal components of the GC population, population variability, mean pairwise GC correlation, temporal sparseness, temporal lossiness, population lossiness, and input variance retained) we used two linear regression normalization techniques: LASSO and RIDGE regression. For Figure 6, LASSO was used to isolate the ‘top’ regressors, while RIDGE was used in Figure 7 to preserve small contributions from regressors. The RIDGE regression method was then used to compare resultant regression slopes (beta coefficients) to changes in task parameters (see Methods on Simulation of cerebellar tasks).

Regressions were performed using the fitrlinear() function in MATLAB, with LASSO selected by using the ‘SpaRSA’ (Sparse Reconstruction by Separable Approximation; Wright et al., 2009) solver, and RIDGE selected with the ‘lbfgs’ (Limited-memory BFGS; Nodecal and Wright 2006) solver techniques. The potential spread of MSE in the models was determined using a K-fold validation technique, with 10 ‘folds’ used, as well as for determining the range of slopes shown in Figures 7, B, C, E, F, H, and I, of which the mean and standard deviation of cross-validation trials are plotted with solid lines and shaded polygons, respectively. Models were selected by choosing the model with the least complex fitting parameters (i.e. the model with the highest Lamba) while still falling within the bounds of the model with the minimized MSE plus the standard error (a standard ‘1SE’ method).

To convey the overall contribution of regressors to the above models of MSE, both the slope (e.g. ‘Beta’) (Fig. 7: B, E, H), and the slope relative to the magnitude of all slopes were used as plotted metrics (Fig. 7: C, F, I).

### Simulation of cerebellar tasks

To simulate the input and output relationship observed in cerebellar and cerebellar-related tasks like vestibulo-ocular reflex adaptation (VOR), interval estimation, and motor-kinematic transformations, we adjusted the inputs and target functions in the task used above to mimic these scenarios. For the VOR-like task (Fig. 7 A-C), the inputs were 10% cosines with a fixed period and amplitude (10Hz, Amplitude range [0, 2]) and the rest were OU processes with taus of 100 and means and standard deviations of 0.5, and 0.2. The target functions were also cosines whose periods and amplitudes were identical to the inputs, but which had phase offsets between 0 and pi to mimic phase-offset VOR tasks.

The interval estimation tasks (Fig. 7 D-F) had standard OU inputs with target functions that were step functions with amplitude ranges from 0 to 1 and intervals that ranged from 0 to 1000 ms, which was the maximal extent of the epoch.

Finally, to simulate the transformation between motor commands and kinematic predictions, we used human EMG as a proxy for a motor command-like input signal to the GCL. 30 muscles from 15 bilateral target muscles were used (Delis et al., 2018; Hilt et al., 2018). The target function was a kinematic trajectory recorded simultaneously with the recordings of EMG used for the study. Although many body parts and coordinate dimensions were recorded of the kinematics, we opted to use the kinematic signal with the largest variance to simplify the experiment to a single target function.

## Acknowledgements

We are grateful to Dr. Pauline M Hilt in the Delis laboratory for use of the EMG and kinematic data modeled in Figure 6. We thank the members of the Person lab for helpful feedback on initial drafts of the manuscript and Drs. Dan Denman and Alon Poleg-Polsky for helpful discussions during the development of the study. This work was supported by NRSA NS113409 to JIG and NS114430, NSF CAREER 1749568 and the Simons Foundation as part of the Simons-Emory International Consortium on Motor control to ALP.

## References

Achen CH (1982) Interpreting and Using Regression. Sage University Paper Series on Quantitative Applications in the Social Sciences, vol. 29.

Albergaria C, Silva NT, Pritchett DL, Carey MR (2018) Locomotor activity modulates associative learning in mouse cerebellum. Nature Neuroscience 21, 725–735. doi:10.1038/s41593-018-0129-x

Albus JS (1971) A theory of cerebellar function. Mathematical Biosciences 10, 25–61. doi:10.1016/0025-5564(71)90051-4

Albus JS (1975) Data Storage in the Cerebellar Model Articulation Controller (CMAC). ASME. J. Dyn. Sys., Meas., Control. 97(3): 228–233.

Armano S, Rossi P, Taglietti V, D’Angelo E (2000) Long-term potentiation of intrinsic excitability at the mossy fiber–granule cell synapse of rat cerebellum. J Neurosci 20:5208–5216, pmid:10884304.

Apps R, Garwicz M (2005) Anatomical and physiological foundations of cerebellar information processing. Nat Rev Neurosci 6:297–311

Bengtsson F, Jorntell H (2009) Sensory transmission in cerebellar granule cells relies on similarly coded mossy fiber inputs. Proceedings of the National Academy of Sciences 106, 2389–2394. doi:10.1073/pnas.0808428106

Billings G, Piasini E, Lőrincz A, Nusser Z, Silver RA (2014) Network Structure within the Cerebellar Input Layer Enables Lossless Sparse Encoding. Neuron 83, 960–974. doi:10.1016/j.neuron.2014.07.020

Buonomano DV, Mauk MD (1994) Neural Network Model of the Cerebellum: Temporal Discrimination and the Timing of Motor Responses. Neural Computation 6, 38–55. doi:10.1162/neco.1994.6.1.38

Calame DJ, Becker MI, Person AL (2021) Associative learning underlies skilled reach adaptation. Biorxiv 2021.12.17.473247

Cayco-Gajic NA, Clopath C, Silver RA (2017) Sparse synaptic connectivity is required for decorrelation and pattern separation in feedforward networks. Nature Communications 8. doi:10.1038/s41467-017-01109-y

Cayco-Gajic NA, Silver RA (2019) Re-evaluating Circuit Mechanisms Underlying Pattern Separation. Neuron 101, 584–602. doi:10.1016/j.neuron.2019.01.044

Chabrol FP, Arenz A, Wiechert MT, Margrie TW, Digregorio DA (2015) Synaptic diversity enables temporal coding of coincident multisensory inputs in single neurons. Nature Neuroscience 18, 718–727. doi:10.1038/nn.3974

Crowley JJ, Fioravante D, Regehr WG (2009) Dynamics of fast and slow inhibition from cerebellar Golgi cells allow flexible control of synaptic integration. Neuron 63:843–853. doi:10.1016/j.neuron.2009.09.004 pmid:19778512

D’Angelo E, De Zeeuw CI (2009) Timing and plasticity in the cerebellum: focus on the granular layer. Trends Neurosci 32:30–40. doi:10.1016/j.tins.2008.09.007 pmid:18977038

De Zeeuw CI, Simpson JI, Hoogenraad CC, Galjart N, Koekkoek SK, Ruigrok TJ (1998) Microcircuitry and function of the inferior olive. Trends Neurosci. 21: 391–400

De Zeeuw CI (2020) Bidirectional learning in upbound and downbound microzones of the cerebellum. Nat Rev Neurosci 22: 92–110

Dean P, Porrill J (2008) Adaptive-filter Models of the Cerebellum: Computational Analysis. The Cerebellum 7, 567–571. doi:10.1007/s12311-008-0067-3

Delis I, Hilt PM, Pozzo T, Panzeri S, Berret B (2018) Deciphering the functional role of spatial and temporal muscle synergies in whole-body movements. Scientific Reports 8. doi:10.1038/s41598-018-26780-z

Hilt PM, Delis I, Pozzo T, Berret B (2018) Space-by-Time Modular Decomposition Effectively Describes Whole-Body Muscle Activity During Upright Reaching in Various Directions. Front Comput Neurosci. 2018;12:20. doi:10.3389/fncom.2018.00020

Dino MR, Schuerger RJ, Liu Y, Slater NT, Mugnaini E (2000) Unipolar brush cell: a potential feedforward excitatory interneuron of the cerebellum. Neuroscience 98:625–636.

Duguid I, Branco T, London M, Chadderton P, Hausser M (2012) Tonic Inhibition Enhances Fidelity of Sensory Information Transmission in the Cerebellar Cortex. The Journal of Neuroscience 32, 11132–11143. doi:10.1523/jneurosci.0460-12.2012

Eccles JC, Ito M, Szentágothai J (1967) The Cerebellum as a Neuronal Machine, Springer, New York (1967)

Eriksson, JL, Robert A (1999) The representation of pure tones and noise in a model of cochlear nucleus neurons. The Journal of the Acoustical Society of America 106, 1865–1879. doi:10.1121/1.427936

Fujita M (1982) Adaptive filter model of the cerebellum. Biological Cybernetics 45, 195–206. doi:10.1007/bf00336192

Gall D, Prestori F, Sola E, D’Errico A, Roussel C, Forti L, Rossi P, D’Angelo E (2005) Intracellular calcium regulation by burst discharge determines bidirectional long-term synaptic plasticity at the cerebellum input stage. J Neurosci 25:4813–4822, doi:10.1523/JNEUROSCI.0410-05.2005, pmid:15888657.

Gao Z, Proietti-Onori M, Lin Z, Ten Brinke MM, Boele HJ, Potters JW, Ruigrok TJ, Hoebeek FE, De Zeeuw CI (2016) Excitatory cerebellar nucleocortical circuit provides internal amplification during associative conditioning. Neuron 89:645–657. 10.1016/j.neuron.2016.01.008

Gilmer JI, Person AL (2017) Morphological Constraints on Cerebellar Granule Cell Combinatorial Diversity. The Journal of Neuroscience 37, 12153–12166. doi:10.1523/jneurosci.0588-17.2017

Gilmer JI, Person AL (2018) Theoretically Sparse, Empirically Dense: New Views on Cerebellar Granule Cells. Trends in Neurosciences 41, 874–877. doi:10.1016/j.tins.2018.09.013

Giovannucci A, Badura A, Deverett B, Najafi F, Pereira TD, Gao Z, Ozden I, Kloth AD, Pnevmatikakis E, Paninski L, De Zeeuw CI, Medina JF, Wang SS-H (2017) Cerebellar granule cells acquire a widespread predictive feedback signal during motor learning. Nature Neuroscience 20, 727–734. doi:10.1038/nn.4531

Guo C, Huson V, Macosko EZ, Regehr WG (2021) Graded heterogeneity of metabotropic signaling underlies a continuum of cell-intrinsic temporal responses in unipolar brush cells. Nat Comm 12:5491

Gurnani H, Silver RA (2021) Multidimensional population activity in an electrically coupled inhibitory circuit in the cerebellar cortex. Neuron 109, 1739–1753.e8. doi:10.1016/j.neuron.2021.03.027

Herculano-Houzel S (2010) Coordinated scaling of cortical and cerebellar numbers of neurons. Front Neuroanat 4:12. doi:10.3389/fnana.2010.00012 pmid:20300467

Herzfeld DJ, Hall NJ, Tringides M, Lisberger, SG (2020) Principles of operation of a cerebellar learning circuit. eLife 9. doi:10.7554/elife.55217

Herzfeld DJ, Kojima Y, Soetedjo R, Shadmehr R (2015) Encoding of action by the Purkinje cells of the cerebellum. Nature 526, 439–442. doi:10.1038/nature15693

Houck BD, Person AL (2014) Cerebellar loops: a review of the nucleocortical pathway. Cerebellum 13:378–385. 10.1007/s12311-013-0543-2

Houck BD, Person AL (2015) Cerebellar premotor output neurons collateralize to innervate the cerebellar cortex. J Comp Neurol 523:2254–2271. 10.1002/cne.23787

Huang CC, Sugino K, Shima Y, Guo C, Bai S, Mensh BD, Nelson SB, Hantman AW (2013) Convergence of pontine and proprioceptive streams onto multimodal cerebellar granule cells. Elife 2:e00400. doi:10.7554/eLife.00400 pmid:23467508

Hull C (2020) Prediction signals in the cerebellum: beyond supervised motor learning. Elife. 9:e54073. doi: 10.7554/eLife.54073. PMID: 32223891; PMCID: PMC7105376.

Ishikawa T, Shimuta M, Häusser, M (2015) Multimodal sensory integration in single cerebellar granule cells in vivo. eLife 4. doi:10.7554/elife.12916

Ito M, Shiida T, Yagi N, Yamamoto M (1974) Visual influence on rabbit horizontal vestibulo-ocular reflex presumably effected via the cerebellar flocculus. Brain Res 65:170 –174.

Izawa J, Criscimagna-Hemminger SE, Shadmehr R (2012) Cerebellar Contributions to Reach Adaptation and Learning Sensory Consequences of Action. The Journal of Neuroscience 32, 4230–4239. doi:10.1523/jneurosci.6353-11.2012

Jakab RL, Hamori J (1988) Quantitative morphology and synaptology of cerebellar glomeruli in the rat. Anatomy and Embryology 179, 81–88. doi:10.1007/bf00305102

Judd EN, Lewis SM, Person AL (2021) Diverse inhibitory projections from the cerebellar interposed nucleus. Elife. 10:e66231. doi: 10.7554/eLife.66231. PMID: 34542410; PMCID: PMC8483738.

Kalmbach BE, Voicu H, Ohyama T, Mauk MD (2011) A Subtraction Mechanism of Temporal Coding in Cerebellar Cortex. The Journal of Neuroscience 31, 2025–2034. doi:10.1523/jneurosci.4212-10.2011

Kanichay RT, Silver RA (2008) Synaptic and Cellular Properties of the Feedforward Inhibitory Circuit within the Input Layer of the Cerebellar Cortex. The Journal of Neuroscience 28, 8955–8967. doi:10.1523/jneurosci.5469-07.2008

Kennedy A, Wayne G, Kaifosh P, Alviña K, Abbott LF, Sawtell NB (2014) A temporal basis for predicting the sensory consequences of motor commands in an electric fish. Nature Neuroscience 17, 416–422. doi:10.1038/nn.3650

Khilkevich A, Halverson HE, Canton-Josh JE, Mauk MD (2016) Links Between Single-Trial Changes and Learning Rate in Eyelid Conditioning. The Cerebellum 15, 112–121. doi:10.1007/s12311-015-0690-8

Kita K, Albergaria C, Machado AS, Carey MR, Müller M, Delvendahl I (2021) GluA4 facilitates cerebellar expansion coding and enables associative memory formation. eLife 10. doi:10.7554/elife.65152

Knogler LD, Markov DA, Dragomir EI, Štih V, Portugues R (2017) Sensorimotor Representations in Cerebellar Granule Cells in Larval Zebrafish Are Dense, Spatially Organized, and Non-temporally Patterned. Current Biology 27, 1288–1302. doi:10.1016/j.cub.2017.03.029

Krauzlis RJ, Lisberger SG (1991) Visual motion commands for pursuit eye movements in the cerebellum. Science 253:568–71

Lanore F, Cayco-Gajic NA, Gurnani H, Coyle D, Silver, RA (2021) Cerebellar granule cell axons support high-dimensional representations. Nature Neuroscience 24, 1142–1150. doi:10.1038/s41593-021-00873-x

Lincoln JS, Mccormick DA, Thompson RF (1982) Ipsilateral cerebellar lesions prevent learning of the classically conditioned nictitating membrane/eyelid response. Brain Research 242, 190–193. doi:10.1016/0006-8993(82)90510-8

Litwin-Kumar A, Harris KD, Axel R, Sompolinsky H, Abbott LF (2017) Optimal Degrees of Synaptic Connectivity. Neuron 93, 1153–1164.e7. doi:10.1016/j.neuron.2017.01.030

Liu, Y, Tiganj, Z, Hasselmo, ME, Howard, MW (2019) A neural microcircuit model for a scalable scale-invariant representation of time. Hippocampus. 29: 260–274.

Mapelli L, Rossi P, Nieus T, D’Angelo E (2009) Tonic activation of GABAB receptors reduces release probability at inhibitory connections in the cerebellar glomerulus. J Neurophysiol 101:3089–3099. doi:10.1152/jn.91190.2008 pmid:19339456

Marr D (1969) A theory of cerebellar cortex. The Journal of Physiology 202, 437–470. doi:10.1113/jphysiol.1969.sp008820

Martin TA, Keating JG, Goodkin HP, Bastian AJ, Thach WT (1996) Throwing while looking through prisms: I. Focal olivocerebellar lesions impair adaptation. Brain 119, 1183–1198. doi:10.1093/brain/119.4.1183

Mauk MD, Buonomano DV (2004) THE NEURAL BASIS OF TEMPORAL PROCESSING. Annual Review of Neuroscience 27, 307–340. doi:10.1146/annurev.neuro.27.070203.144247

Mauk MD, Steinmetz JE, Thompson RF (1986) Classical conditioning using stimulation of the inferior olive as the unconditioned stimulus. Proc Natl Acad Sci USA, 83 pp. 5349–5353

Mccormick DA, Clark GA, Lavond DG, Thompson RF (1982) Initial localization of the memory trace for a basic form of learning. Proceedings of the National Academy of Sciences 79, 2731–2735. doi:10.1073/pnas.79.8.2731

Medina J (2000) Mechanisms of cerebellar learning suggested by eyelid conditioning. Current Opinion in Neurobiology 10, 717–724. doi:10.1016/s0959-4388(00)00154-9

Miall RC, Wolpert DM (1996) Forward Models for Physiological Motor Control. Neural Netw. 9:1265–1279.

Millenson JR, Kehoe EJ, Gormezano I (1977) Classical conditioning of the rabbit’s nictitating membrane response under fixed and mixed CS–US intervals. Learn Motiv, 8 pp. 351–366

Najafi F, Giovannucci A, Wang SS, Medina JF (2014) Coding of stimulus strength via analog calcium signals in Purkinje cell dendrites of awake mice. Elife. 2014;3:e03663. doi:10.7554/eLife.03663

Narain D, Remington ED, De Zeeuw CI (2018) A cerebellar mechanism for learning prior distributions of time intervals. Nat Commun. 9, 469.

Nocedal J, Wright SJ (2006) Numerical Optimization, 2nd ed., New York: Springer.

Palay S, Chan-Palay V (1974) Cerebellar cortex: Cytology and Organization, pp 100–132. Berlin-Heidelberg: Springer-Verlag.

Palkovits M, Magyar P, Szentágothai J (1971) Quantitative histological analysis of the cerebellar cortex in the cat. Brain Research 32, 15–30. doi:10.1016/0006-8993(71)90152-1

Rancz EA, Ishikawa T, Duguid I, Chadderton P, Mahon S, Häusser M (2007) High-fidelity transmission of sensory information by single cerebellar mossy fibre boutons. Nature 450, 1245–1248. doi:10.1038/nature05995

Raymond JL, Lisberger SG (1998) Neural Learning Rules for the Vestibulo-Ocular Reflex. The Journal of Neuroscience 18, 9112–9129. doi:10.1523/jneurosci.18-21-09112.1998

Raymond JL, Medina JF (2018) Computational Principles of Supervised Learning in the Cerebellum. Annu Rev Neurosci. 41:233–253. doi: 10.1146/annurev-neuro-080317-061948. PMID: 29986160; PMCID: PMC6056176.

Rizwan AP, Zhan X, Zamponi GW, Turner RW (2016) Long-Term Potentiation at the Mossy Fiber– Granule Cell Relay Invokes Postsynaptic Second-Messenger Regulation of Kv4 Channels. The Journal of Neuroscience 36, 11196–11207. doi:10.1523/jneurosci.2051-16.2016

Rossi P, D’Angelo E, Taglietti V (1996) Differential long-lasting potentiation of the NMDA and non-NMDA synaptic currents induced by metabotropic and NMDA receptor coactivation in cerebellar granule cells. Eur J Neurosci 8:1182–1189, doi:10.1111/j.1460-9568.1996.tb01286.x, pmid:8752588.

Rudolph S, Hull C, Regehr WG (2015) Active dendrites and differential distribution of calcium channels enable functional compartmentalization of Golgi cells. J Neurosci 35:15492–15504. doi:10.1523/JNEUROSCI.3132-15.2015 pmid:26609148

Ruttle JE, Marius ‘t Hart B, Henriques DYP (2021) Implicit motor learning within three trials. Sci Rep. 11(1):1627. doi: 10.1038/s41598-021-81031-y.

Sanger TD, Yamashita O, Kawato M (2020) Expansion coding and computation in the cerebellum: 50 years after the Marr–Albus codon theory. The Journal of Physiology 598, 913–928. doi:10.1113/jp278745

Saviane C, Silver RA (2006) Fast vesicle reloading and a large pool sustain high bandwidth transmission at a central synapse. Nature 439:983–987. doi:10.1038/nature04509 pmid:16496000

Shadmehr R, Mussa-Ivaldi F (1994) Adaptive representation of dynamics during learning of a motor task. The Journal of Neuroscience 14, 3208–3224. doi:10.1523/jneurosci.14-05-03208.1994

Simat M, Parpan F, Fritschy JM (2007) Heterogeneity of glycinergic and gabaergic interneurons in the granule cell layer of mouse cerebellum. J Comp Neurol 500:71–83. doi:10.1002/cne.21142 pmid:17099896

Smith MA, Ghazizadeh A, Shadmehr R (2006) Interacting adaptive processes with different timescales underlie short-term motor learning. PLoS Biol 4: e179, 2006. doi:10.1371/journal.pbio.0040179

Solinas S, Nieus T, D’Angelo E (2010) A realistic large-scale model of the cerebellum granular layer predicts circuit spatio-temporal filtering properties. Front Cell Neurosci. 2010 May 14;4:12. doi: 10.3389/fncel.2010.00012. PMID: 20508743; PMCID: PMC2876868.

Tabuchi S, Gilmer JI, Purba K, Person AL (2019) Pathway-Specific Drive of Cerebellar Golgi Cells Reveals Integrative Rules of Cortical Inhibition. The Journal of Neuroscience 39, 1169–1181. doi:10.1523/jneurosci.1448-18.2018

Tseng YW, Diedrichsen J, Krakauer JW, Shadmehr R, Bastian AJ. (2007) Sensory prediction errors drive cerebellum-dependent adaptation of reaching. J Neurophysiol. 2007 Jul;98(1):54–62. doi: 10.1152/jn.00266.2007. Epub 2007 May 16. PMID: 17507504.

Tyrrell T, Willshaw D (1992) Cerebellar cortex: its simulation and the relevance of Marr’s theory. Philos Trans R Soc Lond B Biol Sci. 29;336(1277):239–57. doi: 10.1098/rstb.1992.0059. PMID: 1353267.

Wagner MJ, Kim TH, Savall J, Schnitzer MJ, Luo L (2017) Cerebellar granule cells encode the expectation of reward. Nature 544, 96–100. doi:10.1038/nature21726

Wolpert DM, Miall RC, Kawato, M (1998) Internal models in the cerebellum. Trends in Cognitive Sciences 2, 338–347. doi:10.1016/s1364-6613(98)01221-2

Wright SJ, Nowak RD, Figueiredo MAT (2009) Sparse Reconstruction by Separable Approximation. Trans. Sig. Proc., Vol. 57, No 7: 2479–2493.

Yang Y, Lisberger SG (2014) Role of Plasticity at Different Sites across the Time Course of Cerebellar Motor Learning. The Journal of Neuroscience 34, 7077–7090. doi:10.1523/jneurosci.0017-14.2014

Zhou S, Masmanidis SC, Buonomano DV (2020) Neural Sequences as an Optimal Dynamical Regime for the Readout of Time. Neuron. 108(4):651-658.e5. doi: 10.1016/j.neuron.2020.08.020. Epub 2020 Sep 17. PMID: 32946745; PMCID: PMC7825362.

